# Structural basis for regulation of the proteasome 20S core particle by the Parkinsonism-associated proteins FBXO7 and PI31

**DOI:** 10.64898/2026.05.28.728263

**Authors:** Frank Adolf, Ellen A. Goodall, Ekampreet Singh, Susanne von Gronau, Erignacio Fermin Perez, Darlene Fung, Benedikt Müller, Joao A. Paulo, John Hanna, J. Wade Harper, Brenda A. Schulman

## Abstract

FBXO7 and PI31 variants are linked to rare Parkinsonian syndromes, implicating their dysfunction in neurodegeneration. We define how both engage each other and regulate the proteasome 20S core particle (CP). In cells, each can associate independently with the proteasome, with multiple domains in FBXO7 contributing. Utilizing cryo-EM we visualized how FBXO7’s C-terminal domain engages multiple subunits within the CP interior, blocking the β5 peptidase activity. In contrast, we visualized PI31 engagement of all three catalytic sites within the 20S CP, revealing the previously unknown structural basis for β1 inhibition. Furthermore, we establish how disease-associated variants impact both FBXO7 and PI31 function, including disruption of proteasome inhibition and SKP1–FBXO7–PI31 complex assembly. These results establish an unexpected function for FBXO7, providing a mechanistic basis for investigation of its role in proteasome regulation in Parkinson’s disease.

## Introduction

Extensive genetic studies over the last two decades have identified more than two dozen risk variants linked with Parkinson’s disease (PD), which are referred to as PARK genes^1,2^. PARK15 was initially described as a recessive risk factor for Parkinsonian-Pyramidal Syndrome (PPS), a rare juvenile-onset neurological disorder with some phenotypes reminiscent of PD. It is now recognized that PARK15 is encompassed by variants in F-box only protein 7 (FBXO7)^3-12^ (reviewed in refs^13-15^). F-box proteins function as substrate receptor subunits of SCF (SKP1– CUL1–F-box protein) E3 ubiquitin ligases. Through their conserved F-box domain, they bind the adaptor SKP1, which links the F-box protein to Cullin1 (CUL1). Other F-box protein domains mediate regulation, and the recruitment of specific substrates to the core ubiquitin ligase complex^16^.

FBXO7 is a multidomain protein. Alongside its F-box and N-terminal ubiquitin-like (UBL) domain, FBXO7 also contains a central FP domain and a C-terminal proline rich region (PRR). The FP domain takes its name from its presence in FBXO7 and only one other human protein, Proteasome Inhibitor of 31 kDa (PI31; encoded by *PSMF1*). PI31 also contains an intrinsically disordered C-terminal PRR^17^, although whether the PRR domains of FBXO7 and PI31 share structural or functional similarity remains unknown.

FBXO7 and PI31 are capable of forming either homo- or heterodimers via their FP domains^17^. Accordingly, FBXO7 and PI31 reciprocally co-immunoprecipitate in human cells^11,18^, and form a complex when expressed and purified from insect cells^19^. More recently, PI31 variants have been linked to recessive early-onset forms of PD^20^, suggesting that both proteins may function within a shared disease-relevant pathway.

Candidate disease risk alleles map throughout both proteins, suggesting multiple domains may contribute to disease suppression^3,4,13,20,21^. Notably, mutations in FBXO7 are present in the F-box along with additional domains previously implicated in substrate recruitment, supporting the idea that intact E3 ligase activity is required to limit disease progression^22,23^. Also, crosstalk between FBXO7 and PI31 was implied by studies of fibroblasts from patients harboring FBXO7 mutation. These displayed reduced PI31 levels, as well as reduced proteasomal activity and abundance^11^, although such alterations are not observed in induced neuron or HeLa cells carrying engineered FBXO7 deletions^19^.

Multiple lines of evidence link PI31 and FBXO7 with the proteasome. First, PI31 was initially described as an *in vitro* inhibitor of the proteasome 20S core particle (CP)^24,25^, a 28-subunit, barrel-shaped structure composed of four stacked rings each containing seven unique subunits. Two outer α-rings comprise α1-α7 subunits, and two inner β-rings comprise β1-β7 subunits. Three β-subunits are proteases with distinct substrate specificities: β1, caspase-like; β2, trypsin-like; and β5, chymotrypsin-like. The protease active sites are sequestered within the barrel, thereby preventing non-specific protein degradation^26-28^. Access to the proteolytic chamber occurs via a retractable gate present in each α-ring^29^. These gates can be opened by a variety of stimuli, typically via interaction with proteasome caps including the 19S regulatory particle (19S RP)^30-33^, 11S regulators (PA28 family members)^34-36^, and PA200/BLM10^37-39^ (reviewed in refs^40-42^). The α-ring gates can also be artificially opened by deletion of the N-terminal tails of α-subunits^29,30,43^ or chemical approaches (e.g. low concentrations of SDS)^44^. Second, a diversity of functional roles proposed for PI31 converge on proteasome regulation. PI31 has been proposed as: 1) a positive regulator of proteasome activity^45,46^, 2) an adaptor for proteasome trafficking on microtubules^47^, 3) protective of neuronal health with accumulation of polyubiquitin in PSMF1 knockouts^48^ and overexpression ameliorating phenotypes of FBXO7 disruption^49^ as well as 4) a positive or negative modulator of immunoproteasome function^50-52^. Third, both PI31 and FBXO7 co-immunoprecipitate 20S CP subunits (and vice versa)^18,22,53-55^, although it remains unknown if association of FBXO7 with the proteasome requires PI31. In a yeast two-hybrid assay, FBXO7 interacted with the α2 subunit^22^. Meanwhile, deletion of the UBL domain reduced, but did not eliminate, FBXO7 interaction with CP subunits in cells^22^. Electron cryo microscopy (cryo-EM) studies showed PI31 and its orthologs, including budding yeast Fub1 and a microsporidian PI31-like protein, in complex with 20S CP. The structures provided insights into how distinct PI31 PRRs directly engage active sites to inhibit 20S CP activity and prevent PRR domain cleavage^56-58^. Despite strong evidence that PI31 suppresses the β1 caspase-like activity *in vitro*^25,56^, the structural mechanism underlying this inhibition remains unresolved, as human PI31 sequences were not visualized at the β1 active sites^56^, and different primary sequences mediate β1 inhibition in the yeast Fub1 and microsporidian PI31-like structures^57,58^.

Here, we present structural and functional characterization of the interactions between human PI31 and FBXO7, both with each other and with the 20S CP. Utilizing immunoprecipitation combined with quantitative mass spectrometry we demonstrate that human FBXO7 and PI31 can each associate with the 20S CP independently via multivalent interactions mediated by their C-terminal PRRs, as well as FBXO7’s N-terminal UBL-domain. Characterization of a minimal SCF^FBXO7/PI31^ complex reveals the structural basis for FBXO7-PI31 heterodimerization and suggests how PPS-associated variants disrupt this interaction. High-resolution cryo-EM reconstructions of recombinant 20S CP with an engineered open gate allowed visualization and biochemical characterization of how the PRRs of PI31 and FBXO7 each engage the human proteasomal active sites. Our data utilizing model substrates link PD-associated PI31 variants within the PRR to defects in 20S CP interaction. Overall, these data suggest functional interplay between FBXO7 and PI31 in their interaction with 20S CP, providing a mechanistic basis for how disruptions in PI31 and FBXO7 may contribute to PD.

## Results

### Multiple modes of FBXO7-PI31 interaction with the proteasome

To examine whether association of FBXO7 with the proteasome requires PI31, we used Tandem Mass Tagging (TMT)-based proteomics to quantify FBXO7-associated proteins (**Fig. 1a-d, Extended Data Fig. 1a**). Immunoprecipitation was performed for FLAG-tagged FBXO7 expressed with or without untagged full length PI31 or PI31^ΔPRR^ in FBXO7^-/-^; PI31^-/-^ HEK293T cells (**Extended Data Fig. 1a**). As assessed by log2 fold change (FC) relative to negative control FLAG-GFP immunoprecipitates from HEK293T cells, FLAG-FBXO7 immunoprecipitates were primarily enriched in proteasome subunits or components of the SCF (e.g. SKP1 and CUL1) (**Fig. 1b,d, Extended Data Fig. 1b**). In correlation plots of complexes from cells with or without PI31 expression, proteasome and SCF components were present along the diagonal. In the absence of PI31, CSN/signalosome components and 19S RP subunits were relatively more or less enriched, respectively, as compared with 20S CP subunits (**Fig. 1b,d, Extended Data Fig. 1b**). Similarly, in cells expressing PI31^ΔPRR^, FBOX7 retained ability to interact with proteasome and SCF components (**Fig. 1c, Extended Data Fig. 1b**).

**Fig 1:**
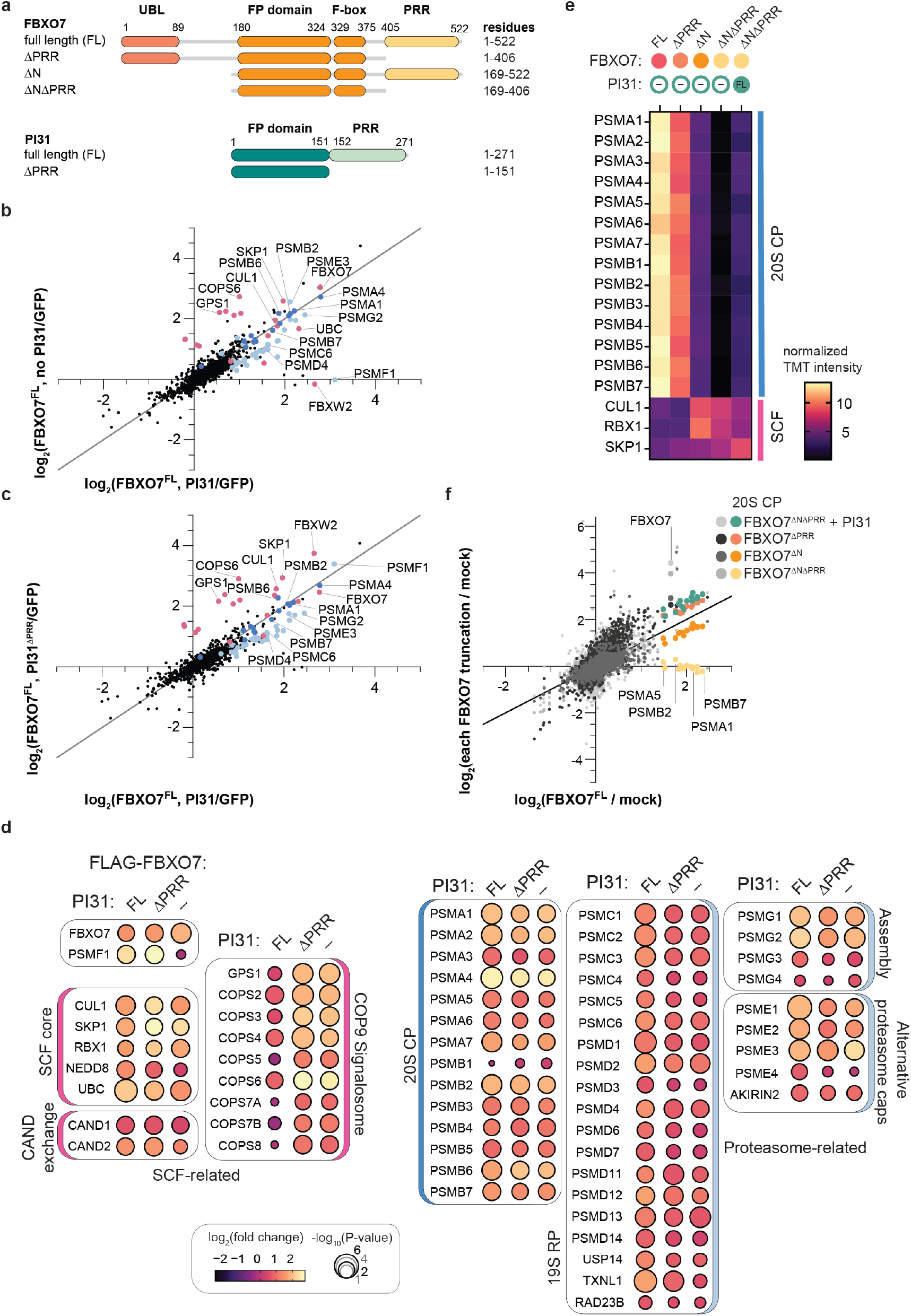
FBXO7–PI31 proteasome interaction. (**a**) Schematic representation of the domain organization of FBXO7 and PI31. UBL, ubiquitin-like; FP, dimerization domain found in FBXO7 and PI31; F-box, domain that associates with SKP1; PRR, proline-rich region. (**b,c**) Correlation plots of log2FC for enriched proteins in FLAG-FBXO7 versus FLAG-GFP control immune complexes from the indicated HEK293T cells (see **Extended Data Fig. 1 a**) with or without PI31 (**b**) or with PI31^ΔPRR^ (**c**) expression. All proteins detected with at least two peptides quantified are plotted as black points. SCF components, COP9 signalosome and CAND exchange factors are highlighted in pink; 20S CP subunits in dark blue; and proteasome-associated components including 19S RP subunits and binding proteins, alternative caps, and proteasome assembly chaperones in light blue. Specific examples from each of these groups are labeled by name on the plots. (**d**) Summary of log2FC and p-values for all proteasome and SCF components identified from the experiment depicted in panels **b** and **c**. (**e**) Heat map displaying the normalized TMT intensities for enriched 20S CP and SCF components measured for the immune complexes from FBXO7^-/-^;PI31^-/-^ HEK293T cells with the indicated FLAG-FBXO7 constructs, with or without co-expression of PI31. (**f**) Overlay of log2FC correlation plots comparing FBXO7^FL^ enrichment over control to the indicated FBXO7 constructs. Color code refers to CP subunits with the indicated FBXO7 construct. Grayscale dots refer to all other proteins detected with at least two peptides quantified.

The extent of enrichment and associated p-values for all detected SCF and proteasome components are summarized in **Fig. 1d**, highlighting 20S CP association with FBXO7 independent of PI31. PI31-independent interaction of FBXO7 with the CP β5 subunit and with CUL1 was validated using immunoblotting (**Extended Data Fig. 1c**).

To define FBXO7 regions mediating proteasome binding, we performed experiments with FLAG-FBXO7 constructs lacking residues 1-168 to remove the UBL (ΔN), or residues 407-522 to remove the PRR domain (ΔPRR), or combining both deletions (ΔNΔPRR) (**Fig. 1a,e,f, Extended Data Fig. 2a, Extended Data Table 2**). As expected, all these FBXO7 constructs associated with core SCF subunits (SKP1, CUL1, RBX1), consistent with their retaining the F-box domain (**Fig. 1a,e**). Moreover, all 20S CP subunits were enriched in FBXO7^FL^ immunoprecipitates from cells lacking PI31, further validating the results described above in an independent experiment. Furthermore, both FBXO7^ΔN^ and FBXO7^ΔPRR^ maintained interaction with CP subunits, albeit at reduced levels, when compared with FBXO7^FL^ (**Fig. 1e,f, Extended Data Fig. 2b-e**). Only with FBXO7^ΔNΔPRR^ was interaction with proteasome subunits lost; however, co-expression of PI31 with FBXO7^ΔNΔPRR^ restored interaction with the CP comparable to the levels observed with FBXO7^ΔN^ (**Fig. 1e,f, Extended Data Fig 2b,c**). These proteomic results were validated using immunoblotting for CP subunit β5 and SCF component CUL1 (**Extended Data Fig 2f**). Taken together, these data suggest multiple potential modes of interaction between FBXO7-PI31 complexes and 20S CP, including interaction of FBXO7 with the proteasome in the absence of PI31 through both its N-terminal UBL domain^22^ and its C-terminal PRR. The data (**Fig. 1e,f**) also suggest that the PRR domain of PI31 can associate with the CP in the context of a complex with FBXO7 lacking both its UBL and PRR domains.

### Structural basis for FBXO7–PI31 heterodimerization within an SCF

To structurally define how FBXO7 engages PI31, SKP1 and CUL1, we obtained a cryo-EM map of a minimal SCF^FBXO7/PI31^ subcomplex, comprising FBXO7 (residues 129-398), PI31 (residues 1-151), SKP1, and CUL1 N-terminal domain (NTD, residues 1-410). The resulting reconstruction, resolved to an overall resolution of 4.2 Å, revealed clear density for a subcomplex comprising the FBXO7 FP–F-box region bound to PI31’s FP domain, and the core SCF core components (SKP1 and the most N-terminal cullin-repeat of CUL1) (**Fig. 2a,b, Extended Data Fig. 3)**. Notably, the cryo-EM data matches an AlphaFold3 prediction (**Extended Data Fig. 4a**). The model captures the expected interactions between components, while also revealing previously unrecognized features. First, as in the previously determined structures of these individual regions, the FP domains of FBXO7^59^ and PI31^17^ adopt homologous folds. These FP domains feature a twisted β-sheet on one face, and helices arranged around a long (6-turn) helix on the other. In the complex, the β-sheet of the FBXO7 FP domain packs against the analogous β-sheet in the PI31 FP domain (**Fig. 2c**). This arrangement matches that in a homodimeric crystal structure of the PI31 FP domain^17^, but not other configurations that were proposed^60^ prior to development of AlphaFold^61,62^. Importantly, the PPS-associated FBXO7 L250P variant that impedes heterodimerization with PI31^11^ places a proline residue within a β-strand at the center of the FP domain interface. The disease-associated PI31 L53M variant^20^, however, lies on the opposite face of the FP domain and is unlikely to directly affect heterodimerization (**Fig. 2c**). Second, the SCF interface matches the canonical SKP1–CUL1–F-box assembly^63^. Two PPS-associated FBXO7 mutation sites map to F-box domain regions that play roles in SCF assembly (**Fig. 2d**).

**Fig. 2:**
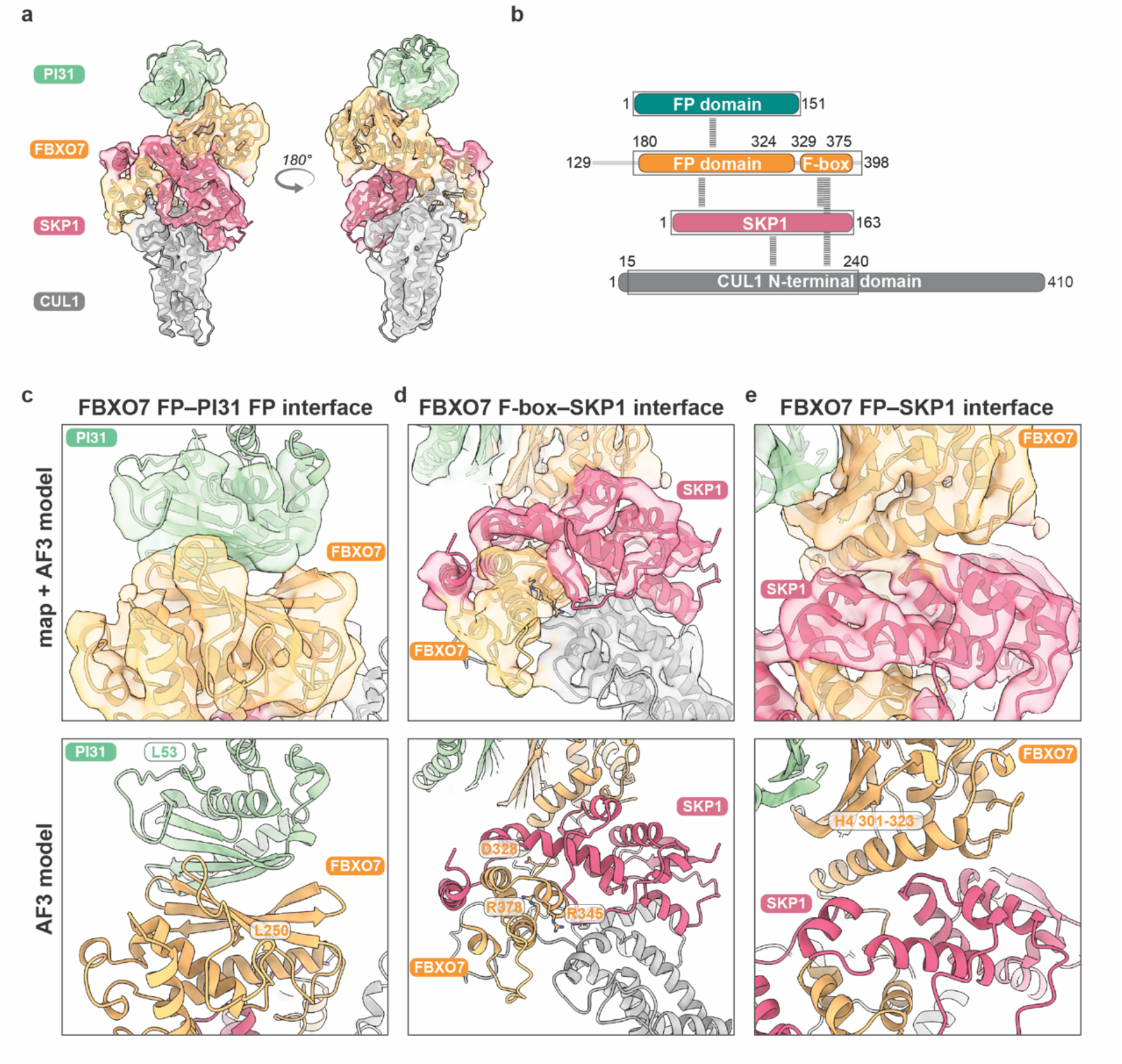
Cryo-EM structure of the SCF^FBXO7–PI31^ complex. (**a**) Overview of the cryo-EM density of the FBXO7–SKP1–PI31–CUL1 N-terminal domain (NTD) complex with fitted AlphaFold3 (AF3) model. PI31 is show in seafoam green, FBXO7 in orange, SKP1 in pink, and CUL1 in gray. (**b**) Schematic representation the complex used for structural determination. Regions resolved in the cryo-EM density are with the modeled portions boxed. Dashed lines indicate domain–domain interfaces. (**c-e**) Inset views of domain–domain interfaces observed in the cryo-EM density. For each interface cryo-EM map with fitted AF3 model (upper panels) and AF3 model alone (lower panels) are shown. (**c**) Inset view of PI31–FBXO7 FP domain interface. Residues mutated in PPS, FBXO7 L250 and PI31 L53 are depicted. (**e**) Inset view of the FBXO7 F-box and SKP1 interface. FBXO7 residues mutated in PPS present in its F-box domain (D328, R345, and R378) are depicted. (**d**) Inset view of FBXO7 FP–SKP1 interface. Helix 4 (H4 301-323) of the FP domain from FBXO7 which packs back against SKP1 is indicated.

The structure also shows extensive interactions between other FBXO7 regions and SKP1. Here, the side of SKP1 opposite its cullin-binding surface is engaged by the linker between the FBXO7 F-box and FP domains, and the FP domain’s long helix (residues 301-323) opposite the PI31-binding β-sheet (**Fig. 2e)**. While this long helix is a central structural feature of FP domains, its sequence is only strongly conserved along the SKP1-binding face in FBXO7 proteins across evolution. Such conservation is not seen in the corresponding helix in PI31, which is exposed in the structure (**Fig. 2a and c, Extended Data Fig. 4c**). Notably, a PPS-associated variant in FBXO7 maps to this noncanonical interface with SKP1 (**Fig. 2d**).

Finally, compared to other SCF assemblies, the FP domains for the FBXO7–PI31 complex are not located in the positions typically observed for substrate-binding domains. Instead, the FBXO7 FP domain is positioned in a location resembling that of the dimerization domain of another SCF complex (SCF^β-TRCP^)^64^ (**Extended Data Fig. 4d**). However, none of the proteasome-interacting domains of FBXO7 or PI31—i.e. FBXO7’s N-terminus, and both C-terminal PRR domains—are present in this assembly, but they are all additional possibilities for substrate-binding domains^22,23^.

### FBXO7 and PI31 differentially inhibit proteasome activities

The sequence and structural similarities between the FP domains of PI31 and FBXO7 have been long recognized^17^. Our finding that FBXO7’s PRR shares functional similarity with that of PI31 in binding to 20S proteasomes (**Fig. 1e,f**) prompted us to further compare and contrast these elements as well. PI31 is well-established as an inhibitor of the 20S CP proteases. Activities of the β1, β2, and β5 proteases are assessed after the 20S CP gates are opened^25,65^. Previous work demonstrated that opening the gates by deletion of the N-terminal tails of the α3 subunit, or of the α7 subunit, permits proper assembly of a 20S CP with constitutively active proteases^29,30,43^. To be able to rigorously compare FBXO7 and PI31 PRRs for their effects on 20S proteasomes, we engineered a recombinant proteasome^66^ with a constitutively open gate (OG) that lacks the N-terminal tails of the α3 and α7 subunits. Cryo-EM analysis confirmed that our recombinant OG-20S CP largely resembles a WT 20S proteasome. Integrity of the α- and β-rings is maintained, while the α-ring gates are open (**Fig. 3a, Extended Data Fig. 5a**). In agreement with previous characterizations^43^, OG-20S CP exhibits elevated basal peptidase activities compared to native 20S CP. Catalytic activity of OG-20S CP was comparable to PA28αβ-activated WT 20S proteasomes (**Fig. 3b**).

**Fig. 3:**
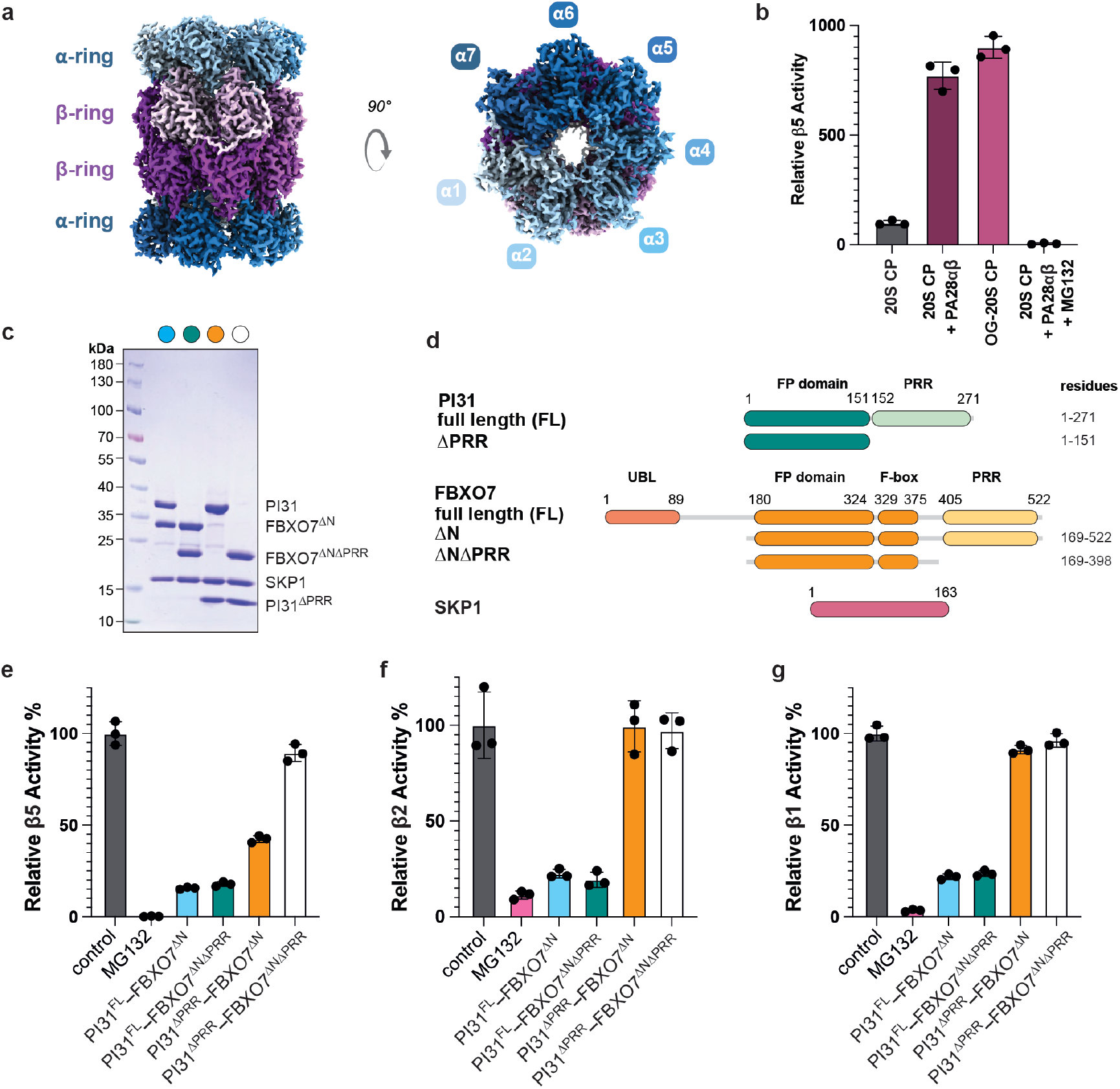
Inhibition of proteasome activity by PI31–FBXO7–SKP1 complexes. (**a**). Cryo-EM density of open gate (OG)-20S CP. The α subunits are colored in shades of blue and β subunits in shades of purple. (**b**) Comparison of β5 chymotrypsin peptidase activity determined by cleavage of suc-LLVY-AMC of recombinant 20S CP alone or in complex with PA28αβ (with or without the inhibitor MG132) to activity of recombinant OG-20S CP. (**c**) Coomassie-stained SDS-PAGE gel of each of the indicated trimeric PI31–FBXO7– SKP1 complexes used in peptidase activity assays (**d**) Domain diagram of PI31, FBXO7 and SKP1 to visualize composition of complex used in peptidase activity assays. (**e-g**) Peptidase activity assays with site-specific fluorogenic peptides for (**e**) β5 chymotrypsin-like activity: suc-LLVY-AMC, (**f**) β2 trypsin-like activity: Z-VVR-AMC, and (**g**) β1 caspase-like activity: Z-LLE-AMC. Protease activity of OG-20S CP (15nM) was assayed in the absence (control) or presence of the indicated PI31–FBXO7–SKP1 trimeric complex (37.5-fold molar excess relative to each class of catalytic active sites) or the pan-proteasome inhibitor MG132 (1.125 µM). Mean activity slopes are plotted, with individual replicates shown. Error bars represent standard deviation (n=3).

With constitutively active 20S CPs in-hand, we next established a system to directly compare inhibitory effects of the FBXO7 and PI31 PRRs present in identical backgrounds. Trimeric SKP1–FBXO7^ΔN^ –PI31 complexes were generated that vary only by the presence or absence of FBXO7 or PI31 PRRs (**Fig. 3c-d**). A 37.5-fold molar excess (inhibitor relative to active site) of the complex containing the PI31 PRR (SKP1– FBXO7^ΔNΔPRR^–PI31) nearly completely inhibited all three proteolytic activities, consistent with prior observations for PI31 alone^56^. Comparable inhibition of 20S CP activities was observed for complexes retaining both FBXO7 and PI31 PRRs (SKP1– FBXO7^ΔN^–PI31). Distinct results were obtained for complexes containing only the FBXO7 PRR (SKP1– FBXO7^ΔN^–PI31^ΔPRR^). The β5 chymotrypsin-like activity was reduced by roughly 50% (**Fig. 3e-g**), but there was no impact on β1 or β2 protease activities of the OG-20S CP in our assay. A homodimeric FBXO7 complex (SKP1–GST-FBXO7^ΔN^) - with two FBXO7 PRRs (**Extended Data Fig. 6a-c**) - did not enhance inhibition beyond that of SKP1–FBXO7^ΔN^–PI31^ΔPRR^.

### Structural basis for inhibition of the human 20S CP by FBXO7

To understand how the FBXO7 PRR interacts with the proteasome, complexes comprising FBXO7^ΔN^–SKP1 and OG-20S CP were analyzed by cryo-EM. The resulting map, at an overall resolution of 1.90 Å, shows density corresponding to residues 427-455 from two FBXO7 molecules symmetrically arranged inside the 20S CP. PRR elements unambiguously enter from opposite sides, as density extends from the antechamber on each end to the central catalytic chamber (**Fig. 4a-c, Extended Data Fig. 7a-d**). The remainder of the FBXO7^ΔN^–SKP1 complex was not resolved, likely due to its flexible tethering to the proteasome-bound PRR.

**Fig. 4:**
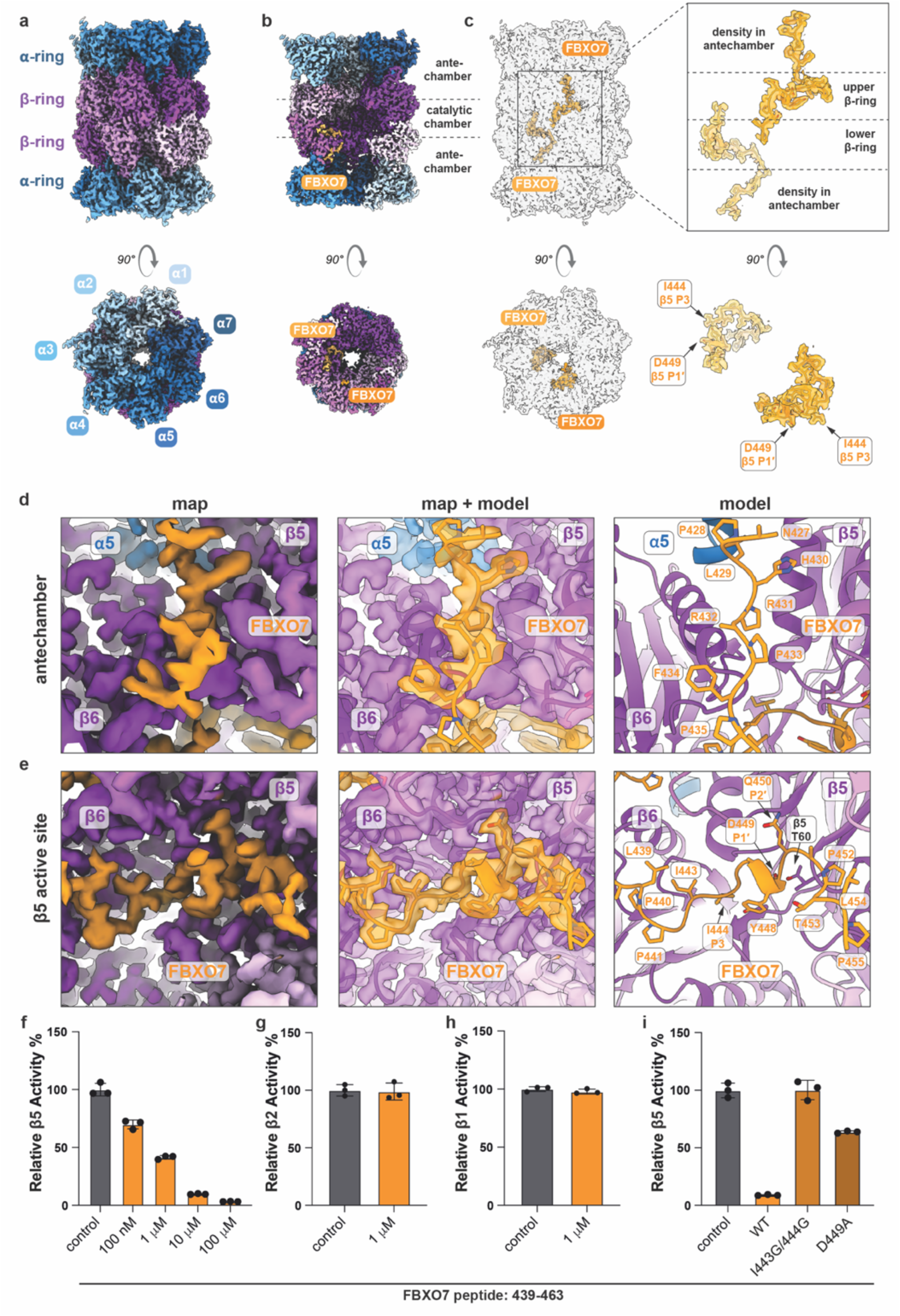
Structure of FBXO7-bound OG-20S CP complex. (**a–c**) Cryo-EM density of the OG-20S CP with segments of the FBXO7 PRR bound within the antechamber and at the β5 active site. Identical side views (top panels) and top views (bottom panels) are shown throughout. 20S CP α subunits are depicted in shades of blue, β subunits in shades of purple, and FBXO7 in shades of orange. **(a)**Overview and (**b**) cutaway views of the cryo-EM density map of the OG-20S CP–FBXO7 complex. The positions of the antechambers and catalytic chamber are indicated. (**c**) The same views as in (a) and (b), with the OG-20S CP rendered transparent to reveal the orientation of two FBXO7 molecules (shades of orange) located in the antechambers and catalytic chamber (left panels). Insets show FBXO7 alone from the same orientations, highlighting PRR segments visualized in the antechamber and catalytic chamber (upper right panel), and residues engaging the β5 active site (lower right panel). (**d–e**) Inset views of FBXO7 PRR bound in antechamber (d) and catalytic chamber (e). Cryo-EM density (left panels), cryo-EM density in transparent with atomic model (middle panels), and the atomic model alone (right panels) are shown. **(d)** Inset view of the interactions of FBXO7 residues 428–435 with β5, β6, and α5 in the antechamber. **(e)** Inset view of the interactions of FBXO7 residues 438–455 within the β5 active site in the catalytic chamber. Residues critical to mimic substrate binding are highlighted. **(f** and **i)** Peptidase activity assays with site-specific fluorogenic peptides for (**f** and **i**) β5 chymotrypsin-like activity: suc-LLVY-AMC, (**g**) β2 trypsin-like activity: Boc-LRR-AMC, and (**h**) β1 caspase-like activity: Z-LLE-AMC. Protease activity of OG-20S CP (5nM) was assayed in the absence (control) or presence of the indicated concentrations of FBXO7 peptide (residues 439-463) (**f–h**) or at 1 µM for comparison of WT and mutant peptides (i). Mean activity slopes are plotted, with individual replicates shown. Error bars represent standard deviation (n=3).

The abundance of proline (27%) and glycine (13%) residues plays critical roles for the FBXO7 PRR. First, it is likely that this region is unstructured in isolation, allowing threading inside the 20S CP. Second, such residues likely enable the extended conformation that outwardly projects proteasome-interacting sidechains such that a small portion of each FBXO7 molecule can span a remarkable end-to-end distance - more than 40 Å through the center of the proteasome barrel. Third, the proline residues punctuate turns that allow the pair of FBXO7 molecules to collectively traverse all three 20S CP chambers (**Fig. 4b,c, Extended Data Fig. 7a-d**) with each PRR directly engaging five distinct subunits (α5, β5, β6, β3, and β4).

Describing the trajectory of each FBXO7 PRR from N-to-C-terminus, the first portion (residues 427-434) embeds in a groove at the α5-β5-β6 interface at the base of the proteasome antechamber. Here, FBXO7 prolines play key roles: Pro428 and Pro431 are located in the antechamber, with Pro428 establishing the trajectory and Pro431 buried in the β5–β6 seam. Pro435 redirects FBXO7 towards the proteasome catalytic chamber (**Fig. 4d**). Residues 436 to 443 enter the catalytic chamber by embracing the β6 surface adjacent to β5. These FBXO7 residues follow a trajectory similar to the β5 propeptide observed in proteasome assembly intermediates^67^. For both FBXO7 and the β5 propeptide, the turn from antechamber to catalytic chamber across β6 is punctuated by a Pro-Gly-Ile motif (FBXO7 residues 441-443), directing the subsequent sequences across the β5 substrate binding groove (**Extended Data Fig. 8c**). Within this groove, conserved residues Ile443, Ile444, Tyr448, and Asp449 of FBXO7 form key contacts. After traversing the β5 surface, FBXO7 is fastened by residues Pro452 to Pro455 at the interface between β4 and the β3–β4 subunits of the opposite β-ring (**Fig. 4e**).

The structure explains how the FBXO7 PRR inhibits β5 activity and evades cleavage itself (**Fig. 4b-e**). Protease cleavage sites in substrates are typically denoted as P3-P2-P1-P1’-P2’-P3’, with the scissile bond between P1 and P1’. Protease binding pockets are correspondingly referred to as S3-S2-S1-S1’-S2’-S3’. Rather than presenting canonical P1 and P1’ residues for cleavage, FBXO7 residues are placed directly in front of and around β5’s active site Thr60. Ile444 is inserted into the β5 S3 pocket, while Asp449 occupies the S1’ position.

The intervening residues, Gly445-Gly446-Glu447-Tyr448, form a small loop, stabilized by hydrogen bonding between Tyr448 of FBXO7 and Thr80 of β5. Densities corresponding to FBXO7 residues Leu439 to Thr453 were well-resolved, surrounding the β5 active site. Notably, we did not observe FBXO7 near the β1 or β2 active sites (**Extended Data Fig. 8a-b**). At present, it remains unknown if the distinct effects of FBXO7 and PI31 towards 20S proteasome activities other than β5 (**Fig. 3d-f**) reflect an inability of FBXO7 to engage the β1 and β2 active sites, or if those interactions occur but are too transient or conformationally heterogeneous to be captured in our cryo-EM reconstructions.

To validate the structural findings, we synthesized an FBXO7 peptide corresponding to this region (residues 439-463). This peptide strongly inhibited the β5 active site in a dose-dependent manner, with a half-maximal inhibitory concentration of <1 micromolar (**Fig. 4f**). It is worth noting that this FBXO7 peptide was significantly more potent than the corresponding peptide from PI31 (IC50 ∼50 micromolar)^68^, which may reflect the more compact inhibitory domain of FBXO7 as described below. Inhibition was specific for β5 function, as the peptide showed little inhibitory effect on cleavage of model substrates for the β2 or β1 proteases (**Fig. 4g-h**). Notably, peptides with substitutions for residues Ile444 (plus Ile443) or Asp449 that occupy the β5 S3 and S1’ pockets, respectively, were defective at inhibition. The I443G/I444G mutant peptide showed complete loss of inhibitory activity, while the D449A peptide showed a strong, albeit partial, loss of inhibition (**Fig. 4i**).

### Structural basis for inhibition of the human 20S CP by PI31

Our structure revealing the basis for β5 inhibition by FBXO7’s PRR prompted a direct comparison with PI31. Previous structures established general modes of PI31 PRR–20S CP engagement, but revealed divergent targeting of β1 in yeast^57^ and microsporidia^58^. Furthermore, in the human PI31–bovine 20S CP complex, interactions were resolved only at the β5 and β2 active sites, leaving the structural basis for β1 inhibition unresolved^56^. To address this, we determined a cryo-EM structure of the human PI31–OG-20S proteasome complex at 2.06Å overall resolution. As in prior structures^56-58^, two PI31 molecules are symmetrically arranged inside the 20S CP (**Fig. 5a-f, Extended Data Fig. 7e-h**). Notably, with well-defined density at the β1 active site, the structure visualizes human PI31 engagement at all three proteasome active sites for the first time. The structure of PI31 at the β5 and β2 active sites (**Fig. 5d, 3**) superimposes with the prior human PI31-bovine 20S proteasome complex^56^. Thus, these structural data validate use of recombinant OG-20S proteasomes, and enable mechanistically defining PI31 inhibition of β1 activity (**Fig. 5f**).

**Fig. 5:**
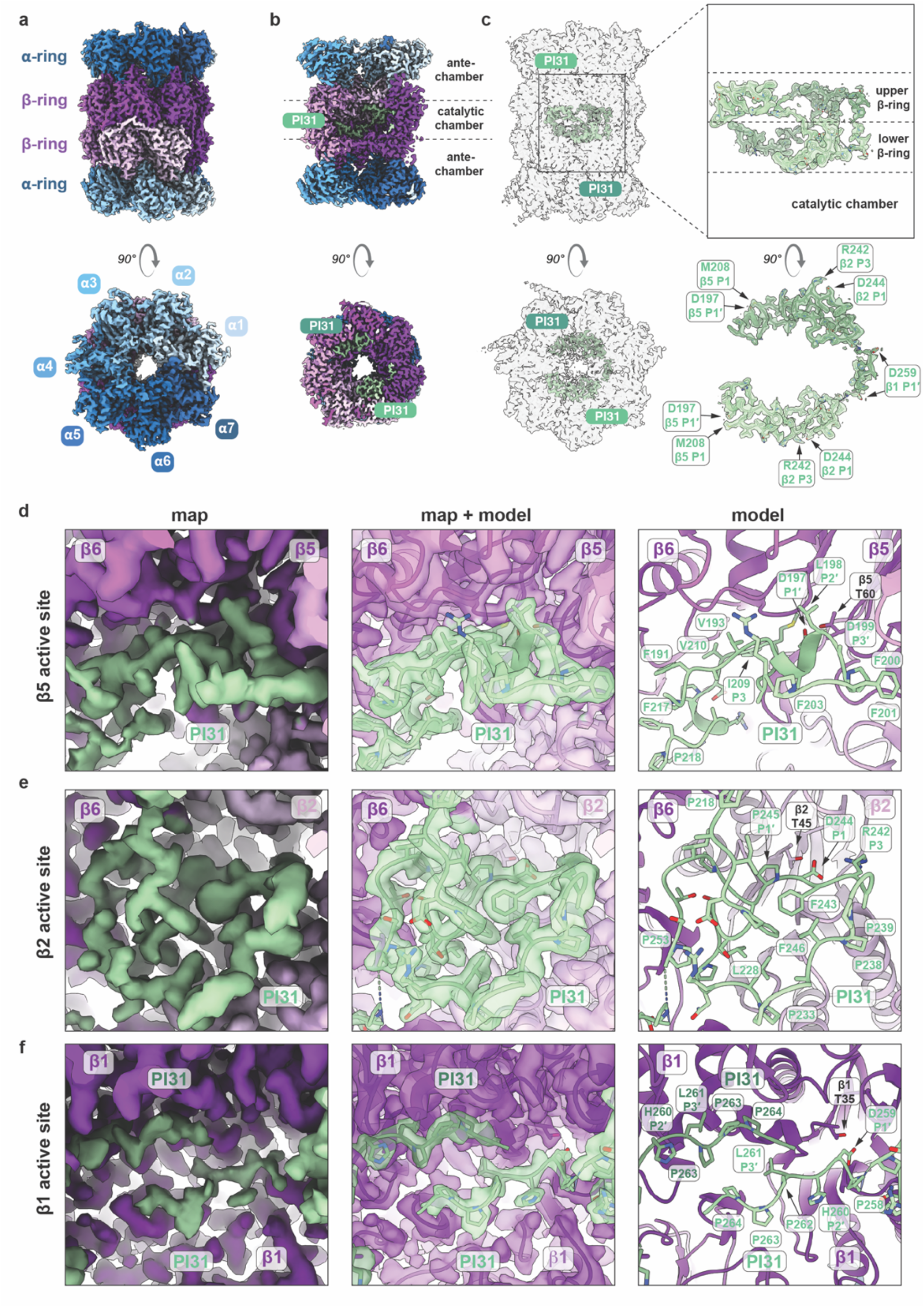
Structure of the PI31 bound OG-20S CP complex. (**a–c**) Cryo-EM density of the OG-20S CP with segments of the PI31 PRR bound in the catalytic chamber engaging with the β5, β2, and β1 active sites. Identical side views (top panels) and top views (bottom panels) are shown throughout. 20S CP α subunits are depicted in shades of blue, β subunits in shades of purple, and PI31 in shades of seafoam green. (**a**) Overview and (**b**) cutaway views of the cryo-EM density map of the OG-20S CP–FBXO7 complex. The positions of the antechambers and catalytic chamber are indicated. (**c**) The same views as in (a and b), with the OG-20S CP rendered transparent to reveal two PI31 molecules (light and medium seafoam green) bound within the catalytic chamber. Insets show PI31 alone from the same orientations, highlighting PRR segments visualized in the catalytic chamber (upper right panel), and residues engaging the β5, β2, and β1 active site (lower right panel). (**d–f**) Inset views of PI31 PRR bound at the (**d**) β5, (**e**) β2, and (**f**) β1 active sites. Cryo-EM density (left panels), transparent density with fitted atomic model (middle panels), and atomic model alone (right panels) are shown. Residues critical to mimic substrate binding are highlighted.

The PI31 segment Pro258 to Pro264 (Pro-Asp-His-Leu-Pro-Pro-Pro) traverses and occupies the β1 active site before terminating at the interface between the β1 and β7 subunits (**Fig. 5f**). The position of this human β1-binding sequence within the PRR, C-terminal to the β2-binding region, is more similar to the microsporidian homolog than the yeast homolog (**Extended Data Fig. 9**). Although the human, microsporidian, and yeast binding modes vary, a common feature of all three structures is the positioning of an aspartate in the S1’ pocket (Asp259 in our structure). The Asp occludes access to the β1 catalytic threonine. The trajectory of the subsequent PI31 residues deviate substantially from a canonical substrate conformation, through extensive stabilizing interactions mediated via the side chains of His260, Leu261, Pro263, and Pro264, all organized by Pro262. These interactions highlight the importance of proline residues in this sequence (**Fig. 5f**).

### Shared and distinct modes of FBXO7 and PI31 PRR domain engagement of proteasome

Comparing human PI31 and FBXO7 structures around the β5 site shows how two different PRR-domain containing proteins can engage the proteasome, inhibit substrate cleavage, and avert their own degradation. Both PI31 and FBXO7 place an aspartate side-chain in the protease oxyanion hole to block the β5 active site Thr60. In both PI31 and FBXO7, the segments engaging the β5 subunit also make extensive contacts with neighboring β6 and β4 subunits within the same β-ring. However, the elements mediating these adjacent interactions differ, as do their trajectories across the β5 active site (**Fig. 6a-b**). FBXO7 acts as a hook as it hugs β6 turning from the antechamber to the catalytic chamber, proceeding from β6 to β5 and the β5 active site threonine. Here, FBXO7 residues Ile444 and Asp449 mimic substrate P3 and P1′ positions as the peptide threads from N-to-C across the β5 active site towards the β4 subunit. By contrast, PI31’s open infinity sign-like shape places all resolved residues within the catalytic chamber (**Fig. 6a-c**). PI31 traverses the β5 site from the opposite direction as FBXO7, first contacting β5 with Asp197 in the S1′ pocket, followed by Leu198 and Asp199 mimicking P2′ and P3′ positions as the sequence traverses to β4. PI31 then loops back across the β5 site to position Met208 (P1) and Ile209 (P3) in the substrate-binding groove, with these residues in the opposite orientation as a substrate would bind.

**Fig. 6:**
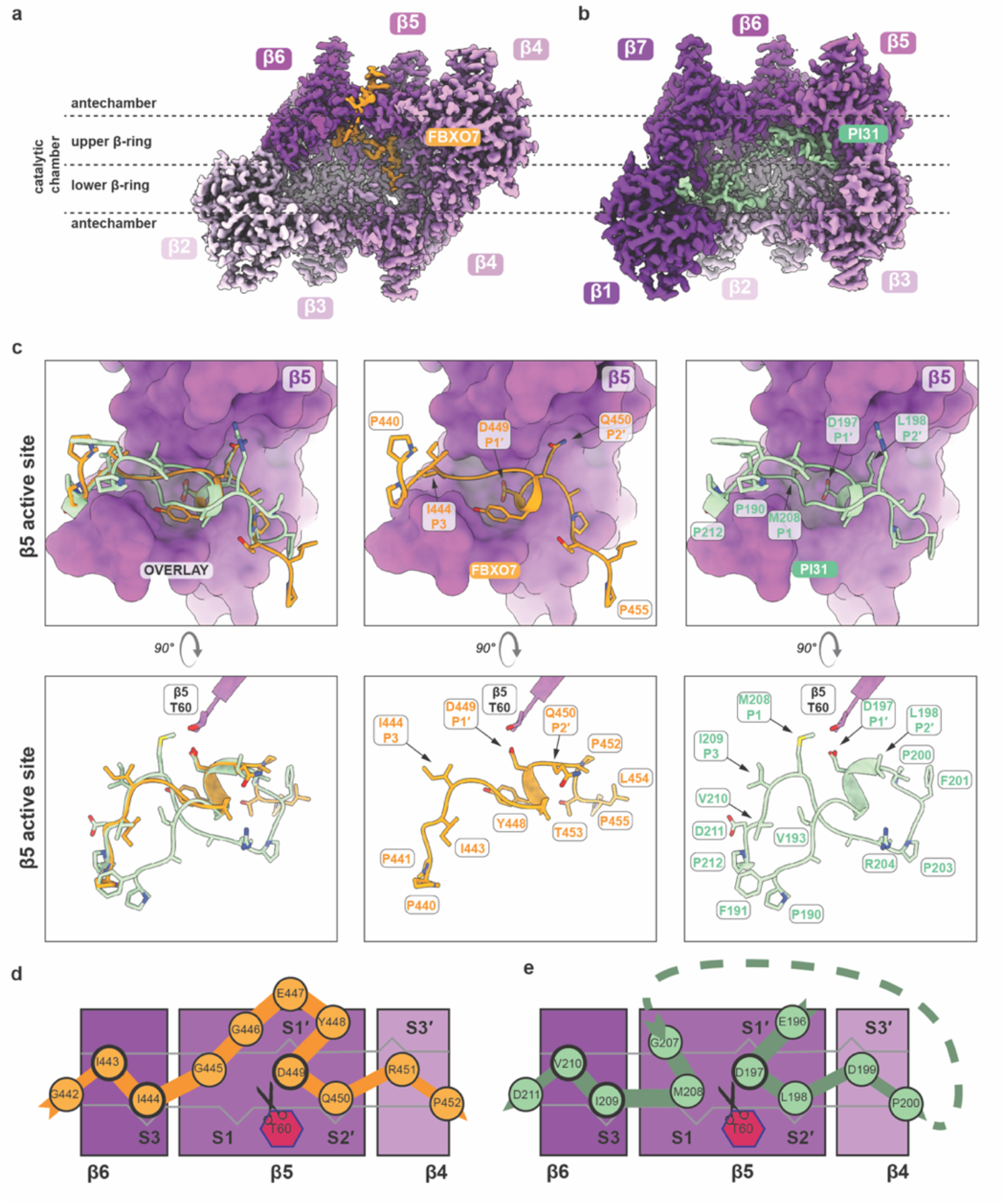
Comparison of FBXO7 and PI31 binding to the 20S CP. (**a–b**) Cryo-EM densities of the FBXO7-bound (a) and PI31-bound OG-20S CP complexes. (**a**) Densities for β6, β5, and β4 of the upper β-ring and β2, β3, and β4 of the lower β-ring are shown together with FBXO7 density. (**b**) Densities for β7, β6, and β5 of the upper β-ring and β1, β2, and β3 of the lower β-ring are shown together with PI31 density. The positions of the antechambers and catalytic chamber are indicated. (**c**) Comparison of FBXO7 and PI31 binding at the β5 active site. Overlay of FBXO7- and PI31-bound at the β5 sites (left panels), FBXO7 bound to the β5 site (middle panels), and PI31 bound to the β5 site (right panels). In the upper panels, the β5 subunit is shown in surface representation and FBXO7 and PI31 are shown in ribbon representation. In lower panels views rotated by 90° are shown to highlight the trajectories of FBXO7 and PI31 within the β5 active site. For clarity, only the catalytic β5 Thr60 residue is shown. Residues mediating substrate-mimetic interactions are indicated throughout. (**d–e**) Schematic representation of how (**d**) FBXO7 and (**e**) PI31 engage the β5 active site through substrate-mimetic binding. Both proteins are proposed to prevent their own degradation through a similar “looping-out” mechanism that positions the polypeptide chain outside a cleavage-competent conformation.

FBXO7 and PI31 employ some shared features to avert their own cleavage. Gly residues loop the polypeptide away from the active site to ensure no peptide bond is placed within reach of the active site threonine. As described above, an aspartate occupies the S1′ pocket. And branched chain amino acids sit within the substrate binding groove of β6 with an isoleucine in the S3 pocket. However, these shared features are differentially achieved by FBXO7 and PI31. They traverse the β6 and β5 portion of the substrate binding groove with reversed orientations (**Fig. 6c-d**).

### Impact of PSMF1 (PI31) PD-associated variants on proteasome binding and inhibition

The ability of PI31 containing its C-terminal PRR to promote recruitment of FBXO7 lacking its UBL and PRR, coupled with structural data on PI31’s interaction with CP, led us to examine the ability of PI31 PD-associated patient variants^20^ to interact with the proteasome in cells. Within PI31’s PRR, variants are clustered around the conserved β2-inhibiting residues with three different substitutions of the key Arg242 to either cysteine, glycine, or histidine (**Fig. 7a**). Arg242 mimics a substrate’s P3 residue in binding to the β2 site, at the interface of the β2 and β3 subunits (**Fig. 5e**). L228V is located at the interface between β1 and β2 potentially helping to position Phe246 in PI31 in the S3′ pocket of the β2 site and organize the overall positioning of residues (**Fig. 5e**). The nonsense substitution PI31^R231X^ removes all the residues responsible for PI31 binding at both β2 and β1 sites, but leaves the β5 site intact (**Fig. 5, Fig. 7a**).

**Fig. 7:**
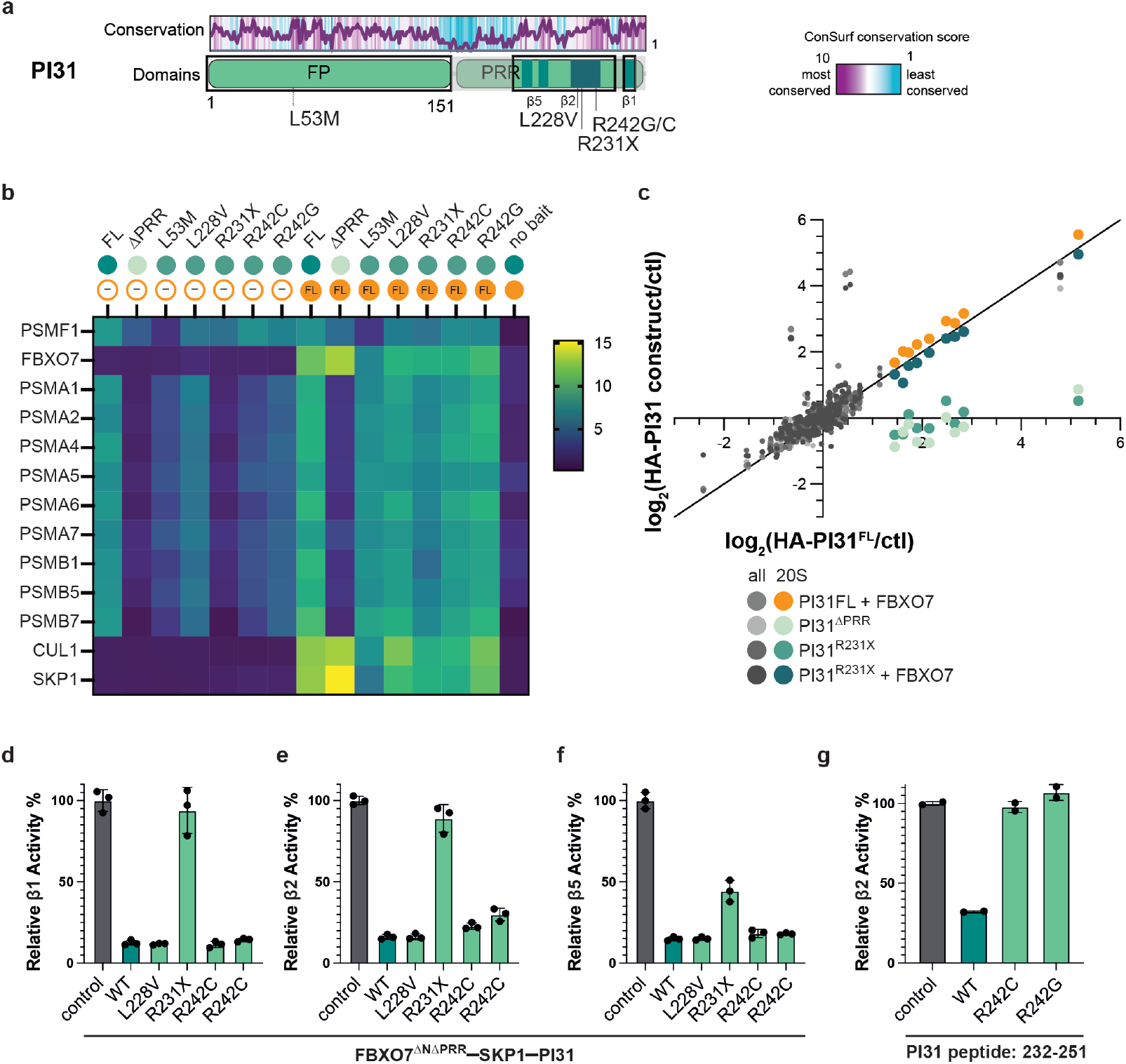
Mapping PD-associated variants in PI31 function to proteasome interaction. (**a**) Domain diagram of PI31 with sequence conservation shown above and sites of PD-associated sequence variants indicated. The sequences resolved are boxed and highlighted. (**b**) Heatmap of relative TMT intensities of 20S CP and SCF components detected for immunoprecipitations for the given PI31 constructs with or without FBXO7. (**c**) Comparison of proteins enriched in HA-tagged PI31 constructs in comparison to those enriched in HA-PI31^FL^ without FBXO7, with enrichment over an untagged control. 20S CP subunits are highlighted in orange (PI31^FL^ + FBXO7), light seafoam (PI31^ΔPRR^), medium seafoam (PI31^R231X^) and dark seafoam (PI31^R231X^ + FBXO7). Each dot detected corresponds to a protein detected with at least three peptides in at least two of three replicate samples. (**d–f**) Peptidase activity assays with site-specific fluorogenic peptides for (**d**) β1 caspase-like activity: Z-LLE-AMC, (**e**) β2 trypsin-like activity: Z-VVR-AMC, and (**f**) β5 chymotrypsin-like activity: suc-LLVY-AMC. Protease activity of OG-20S CP (15 nM) was assayed in the absence (control) or presence of trimeric FBXO7^ΔNΔPRR^–SKP1–PI31 complexes containing the indicated PI31 variant (1.125 µM). Mean activity slopes are plotted, with individual replicates shown. Error bars represent standard deviation (n = 3). **(g)** Peptidase activity assays of the β2 trypsin-like activity assay. Protease activity of native human 20S CP was assayed in the absence (control) or presence of peptides corresponding to PI31 residues 232–251 containing the native sequence or the PD-associated variants R242C or R242G. Mean activity slopes are plotted, with individual replicates shown. Error bars represent standard deviation (n = 2).

To determine the impact of PD-associated variants on PI31, we performed immunoprecipitations of HA-tagged PI31 from FBXO7^-/-^; PI31^-/-^ HEK293T cells and quantified PI31-associated proteins using TMT-based proteomics (**Extended Data Fig. 10a**). First, as expected, HA-PI31 associated with CP subunits in the absence of FBXO7, and this interaction was lost upon removal of its PRR (**Fig. 7b,c**). Second, association of HA-PI31 with SKP1 and CUL1 was dependent on co-expression of FBXO7 (**Fig. 7b**). Third, among the PI31 variants examined in the absence of FBXO7, the R231X variant resulted in the greatest loss in CP interactions, similar to that seen with PI31^ΔPRR^, but importantly, CP interaction was rescued upon co-expression of FBXO7 (**Fig. 7b,c; Extended Data Fig. 10b-e**). Other variants tested (L53M, R242C, R242G, and L228V) displayed a degree of proteasome interaction comparable to wild type PI31 in the presence or absence of FBXO7 expression (**Fig. 7b, Extended Data Fig.10d,c**) The FP-localized variant, PI31^L53M^ was consistently expressed at lower levels, suggesting that this mutation may be destabilizing to the protein overall (**Fig. 7b,c, Extended Data Fig. 10f**).

While proteasome interaction was not abolished for many of the PD-associated variants within PI31’s PRR, we sought to test their impact on peptidase activity. Using FBXO7^ΔNΔPRR^–SKP1–PI31 complexes, we evaluated the impact of the PD-associated variants on binding at each active site locally, as opposed to overall binding to 20S CP. Consistent with the complete removal of residues involved in binding at the β1 and β2 sites, complexes containing PI31^R231X^ were unable to inhibit activity at these sites (**Fig. 7d,e**). At the β5 site, complexes containing PI31^R231X^ remained inhibitory, but were weaker than complexes containing PI31^FL^, perhaps due to a decreased affinity for 20S CP without the remaining residues in PI31 present (**Fig. 7f**). All other PD-associated variants tested, PI31^L228V^, PI31^R242C^ and PI31R^242G^, maintained inhibition at each active site. A peptide corresponding to PI31 residues 232-251 is sufficient to selectively inhibit the β2 activity of 20S CP^68^. We therefore tested the impact of R242C and R242G variants in the context of this peptide. Unlike in the context of full length PI31 PRR, peptides harboring Arg242 variants abrogated inhibition of β2 activity comparable to R242H^68^, a variant also previously identified as an Alzheimer’s Disease risk allele^20,69^ (**Fig. 7g**). These data suggest avidity contributed by multiple 20S CP binding sites within the PRR can be unmasked with the use of peptides sufficient to inhibit individual active sites.

## Discussion

Our study defines structural mechanisms by which the PD-associated proteins FBXO7 and PI31 engage and inhibit the proteasome. First, FBXO7 and PI31 can independently associate with the 20S CP, while also forming an SCF E3 ligase complex (**Fig. 1-2**). Second, the PRRs of both proteins directly engage proteasomal active sites through related but distinct structural mechanisms (**Fig.4-6**). Whereas a single PI31 PRR,inhibits all three proteolytic active sites, we observed inhibition by FBXO7 only of the β5 chymotrypsin-like activity (**Fig. 3**). We additionally resolve the structural basis for human PI31-mediated β1 inhibition, an unresolved question from previous work^56^. Finally, localization of disease-associated variants establishes mechanistic explanations for how some PPS-linked mutations disrupt either SCF assembly or proteasome interaction (**Fig. 2**). Together, these findings reveal that FBXO7 and PI31 are more closely related than previously appreciated, converging on direct regulation of the proteasome, including within a shared SCF E3 ligase complex. In addition, our studies benefited from generation of a recombinant proteasome core particle with constitutively open gates. We note that this reagent could be a useful tool for studying 20S CP activity and for inhibitor development.

Which proteasome populations are targeted by FBXO7 and PI31? Cellular 20S CP abundance exceeds that of FBXO7 and PI31 by at least an order of magnitude^70^, arguing against a model in which these proteins occupy large fractions of the total proteasome pool. Instead, the comparatively low abundance of FBXO7 and PI31 suggests engagement of either a small fraction or specialized forms of the proteasome. Notably, a constitutively open-gate 20S CP mutant produced through the incorporation of an additional copy of α4 upon deletion of α3 from yeast co-purifies with endogenous Fub1, the yeast ortholog of PI31 (ref^57^), much like our recombinant human open gate 20S CP readily binds FBXO7 and PI31. These findings raise the possibility that PI31 and FBXO7 preferentially recognize non-canonical proteasome states with unusually accessible catalytic chambers such as a similar α4-α4 proteasome produced in human cells when relative levels of α4 and α3 subunits are modified^71^.

One possibility is that FBXO7 and PI31 could act as surveillance or quality-control factors for misassembled proteasomes with open gates despite active proteases. Although proteasome biogenesis is highly coordinated to avert such intracellular proteolytic activity, it is possible that some proteasomes could fall through the cracks, for example by misincorporation of highly homologous subunits, or through failure in gate closure during completion of the maturation process. During 20S CP biogenesis, the assembly chaperone complex, PAC1–PAC2 (PSMG1–PSMG2), binds to the α-ring and to POMP within the assembling β-ring, thereby stabilizing an open-gate state^66^. Dissociation of PAC1– PAC2 following autocatalytic cleavage of assembly factors is thought to permit maturation of the closed gate. Meanwhile, after assembly, the gate of fully mature 20S CP is opened through association with regulatory particles, enabling protein degradation. However, incomplete dissociation or failures in gate closure after release of PAC1–PAC2 during the assembly process, or defects in regulatory particle association or dissociation could infrequently generate aberrant open-gate proteasomes. We speculate that FBXO7 and PI31 recognize and possibly, through SCF E3 ligase activity, target these proteasomes. Potential consequences could be directing proteasomes for degradation, influencing proteasome assembly, or directing intracellular trafficking, processes previously linked to both PI31 and FBXO7 (refs.^45,47,51,52^). Interestingly, we find FBXO7 co-immunoprecipitates with PAC1 and PAC2, as well as regulatory particle subunits (**Fig. 1d**). One explanation could be simultaneous proteasome engagement by FBXO7– PI31–SKP1 complex and other proteasome-associated factors to opposite sides of the 20S CP. Visualization of such partially engaged proteasomes would require future studies of complexes assembled under different conditions than ours which relied on excess FBXO7 or PI31.

The structural data suggest that simultaneous binding of the SCF^FBXO7–PI31^ and a canonical regulatory particle to the same α-ring face is sterically incompatible for two reasons. First, the distance that could be spanned by the number of residues between the proteasome-engaging regions of FBXO7 or PI31 and the folded F-box and FP domains is shorter than the distance required to extend between the central catalytic chamber of the proteasome and a regulatory particle entry pore. Second, PRR engagement of the proteasome would place the F-box and FP domains immediately outside the 20S CP, so they would clash with a regulatory particle.

PI31 is required for synaptic proteostasis in vivo, and its loss in mice leads to accumulation of polyubiquitinated proteins, synaptic dysfunction, and progressive neuronal degeneration^48^. Conversely, increased PI31 levels have been shown to suppresses neurodegeneration and extending survival in FBXO7-deficient mouse models^49^. Together, these studies establish a critical role for the PI31–FBXO7 axis in maintaining proteostasis in the nervous system. One challenge in understanding FBXO7 has been discerning the effects of disease-associated variants, which map throughout the entire protein^3-12^, including the FP–FP and F-box–SKP1 interfaces, as well as the proteasome interacting regions (**Extended Data Fig. 4f**). One such variant present in the FP domain of FBXO7 was shown to disrupt binding to PI31 previously^11^, which can be now explained by its mapping to the interface between these proteins in our cryo-EM structure (**Fig. 2**).

As for PD-associated variants within in the PRRs, our data highlight how defects may occur locally, for example in 20S CP active site engagement and inhibition, yet these perturbations may be challenging to detect in the context of multivalent interactions. The relative similarity of PI31 and FBXO7, and their forming a complex with each other as well as the 20S CP, poses a particular challenge in deconvoluting the biochemical and structural basis for mutational defects.

Nonetheless, some mutations clearly impact interplay with the 20S CP. Co-immunoprecipitation experiments suggest an interplay of FBXO7 and PI31 on their interactions with the 20S CP. In one example, a PD-associated PI31 variant which truncates β1- and β2-interacting residues only alters proteasome binding in the absence of FBXO7 (**Fig. 7**). Likewise, PI31 variants disrupting β2-site interactions retain overall proteasome association yet display impaired inhibition when examined using minimal inhibitory peptides. Together these observations illustrate how interactions across multiple binding sites can obscure functionally important local perturbations. The PPS-associated FBXO7 variant R481C^5^ maps to a region analogous to the PI31 β2-inhibitory segment^17^, where multiple PD-associated substitutions of R242 disrupt β2 inhibition (**Fig. 7**). These observations raise the possibility that FBXO7 engages additional active sites under specific conformational or compositional contexts, and such interactions could be impaired by risk alleles.

Overall, our data establish a mechanistic framework for understanding FBXO7–PI31–proteasome interactions and the impact of PD-associated mutations. Our work establishes an unexpected function for FBXO7 in direct proteasome regulation and suggests that impaired proteasome inhibition contributes to Parkinsonian neurodegeneration. Future studies are required to define the physiological contexts and consequences of SCF^FBXO7–PI31^ proteasomes recognition, and how disease-associated variants disrupt this pathway.

## Methods

### Human cell culture

HEK293T cells were cultured in Dulbecco’s modified Eagle’s medium (DMEM), high glucose, pyruvate (Thermo: cat#: 11995065), supplemented with 10% fetal bovine serum (FBS) (Cytiva, cat#: SH30910.03) and penicillin-streptomycin (100 U/mL final concentration) (Thermo cat#: 15140122). Cells were maintained at 37°C with 5% CO2.

### Plasmids and molecular cloning for mammalian expression of FBXO7 and PI31

For generation of lentiviral expression constructs of FBXO7 and PSMF1 in human cell culture, pDONR223 plasmids containing FBXO7 (Addgene # 251201) or PSMF1 (Addgene # 251202) open reading frame and a stop codon were transferred to pDEST vectors: pHAGE N-FLAG-HA, PuroR^72^ (Addgene # 251190);pHAGE pEF1α tagless vector, BlastR (Stephen Elledge); or pLEX305 N-dTAG-HA^73^ (Addgene # 91797), PuroR with LR Clonase (Invitrogen cat#: 11791100) using the Gateway cloning system. Point mutations and truncations in FBXO7 or PSMF1 were incorporated using QuikChange XL mutagenesis (Agilent cat#: 200517) or Q5 site directed mutagenesis (NEB, <u>dx.doi.org/10.17504/protocols.io.bddfi23n</u>), Resulting in vectors encoding FLAG-HA tagged FBXO7 constructs including N and C-terminal truncations (Addgene #s 251177, 251178, 251188,251189), tagless FBXO7 (Addgene # 251191) tagless PI31, both full length (Addgene # 251192) and C-terminally truncated (Addgene # 251193), and dTAG-HA-tagged PI31 constructs including both truncations and point mutants (Addgene #s 251194 -251200).

To generate PSMF1 knockout cells, Cas9 and single guide RNAs (sgRNAs) (PSMF1 guide 1: 5’-CCTTGTGAAAGCCATCACCG-3’ (Addgene # 251203), PSMF1 guide 2: 5’-CCGGTATGAGTATAAGGATG-3’ (Addgene # 251204))were introduced to HEK293T cells previously edited to disrupt FBXO7 using similar methods^19^. PSMF1-targeted guide sequences were introduced into pX458 (Addgene # 48138)^74^. Cells were transfected with these constructs using PEI Max (Polysciences, CAS: 49553-93-7). After three days, GFP expressing cells were single-cell sorted into 96-well plates with a SONY SH800S sorter (RRID:SCR_018066). Selection of clones with gene editing at the PSMF1 locus was confirmed by amplifying the region of PSMF1 targeted by the guides with amplicon primers for each clone (primer sequence: upstream primer 5’-TCCCAATGATAAGAAGTCAGAAC-3’, downstream primer 5’-CTACATGAACATTGAAGCTCTG-3’), next generation sequencing of these amplicons with Illumina Miseq, and analysis with OutKnocker ^75^ and with immunoblotting.

### Stable expression of FBXO7 and PI31 in human cell culture

HEK 293T were used to produce lentiviral particles, where these cells were co-transfected with lentiviral expression vectors along with PMD2.G (Addgene # 12259, from Didier Trono) and psPAX2 (Addgene #12260) using PEI Max (Polysciences, CAS: 49553-93-7) at a ratio of 3:1 PEI to total DNA. One day following transfection, media on cells was exchanged and two days following transfection, media supernatants were harvested from the HEK293T cells as lentiviral supernatant, filtered through 0.45 µ PES syringe filters and applied to cells to be transduced along with 1 µg/mL polybrene. One day following treatment with lentiviral supernatants, media was exchanged for fresh DMEM supplemented with 10% FBS. Two days following lentiviral transduction, cells were treated with 1 µg/mL puromycin for cells treated with pHAGE-FLAG-HA FBXO7 or pLEX305 dTAG-HA-PI31 lentiviral constructs, or 10 µg/mL Blasticidin S for pEF1a-FBXO7 or pEF1a-PI31 constructs with fresh media changes containing selection treatments every alternate day. Cells were then scaled up to triplicate and larger plate format, maintaining antibiotic selections until plating cells for immunoprecipitations.

### FLAG-FBXO7 immunoprecipitations from human cell culture

HEK293T FBXO7^-/-^ PSMF1^-/-^ cells expressing FLAG-HA-FBXO7 constructs were seeded into 150 mm plates with one plate per replicate and performed in triplicate. Once plates reached confluency, or approximately 20 x 10^6^ cells per replicate, cells were scraped in ice cold phosphate buffered saline, pelleted at 500 x g and snap frozen in liquid nitrogen. To begin the immunoprecipitation, all cell pellets were thawed on ice and each mixed end over end with lysis buffer (50 mM HEPES pH 7.6, 140 mM NaCl, 0.5% IGEPAL-CA630 (v/v), protease inhibitors: 1 mM PMSF, 10 µM leupeptin, 1.5 µM pepstatin A, 5 µg/mL aprotinin and Benzonase nuclease (Novagen)) for 30 minutes at 4°C. Lysates were clarified by centrifugation at 21 000 x g for 30 minutes at 4°C followed by total protein quantification using Pierce BCA (cat#: 23250). Normalized protein inputs were incubated with 20 µL of prewashed anti-FLAG M2 magnetic beads (Sigma cat#: M8823) for 3 hours, mixing on orbital plate mixer at 800 rpm at 4°C. Using a Kingfisher Flex (Thermo, RRID:SCR_027576) samples were washed first with 500 µL with low detergent buffer (50 mM HEPES pH 7.6, 140 mM NaCl, 0.1% IGEPAL-CA630 (v/v)),followed by three washes, each of 500 µL of wash buffer (50 mM HEPES pH 7.6, 140 mM NaCl). Finally, beads were eluted with mixing in 75 µL of elution buffer (50 mM HEPES pH 7.6, 140 mM NaCl 0.15 mg/mL,0.15 mg/mL FLAG peptide). Samples from inputs, flowthrough samples, and elutions were set aside for immunoblotting analysis. Eluates were then snap frozen in liquid nitrogen and stored at -80°C until processed for quantitative mass spectrometry.

### HA-PI31 immunoprecipitation from human cell culture

Similar to FLAG-immunoprecipitations, HEK293T FBXO7^-/-^ PSMF1^-/-^ cells expressing dTAG-HA-PI31 constructs were seeded into 6-well plates with one well per replicate, and performed in triplicate. Once plates reached confluency, or approximately 1.2 x 10^6^ cells per replicate, cells were scraped in ice cold phosphate buffered saline, pelleted at 500 x g, snap frozen in liquid nitrogen, and stored at -80°C until continuing with immunoprecipitations., All cell pellets were thawed on ice and each mixed end over end with lysis buffer for 30 minutes at 4°C. Lysates were clarified by centrifugation at 21 000 x g for 30 minutes at 4°C followed by total protein quantification using Pierce BCA (cat#: 23250). Normalized protein inputs were incubated with 9 µL of prewashed anti-HA magnetic beads (Sigma cat#: M8823) for 3 hours, mixing on orbital plate mixer at 800 rpm at 4°C. Using a Kingfisher Flex (Thermo, RRID:SCR_027576) samples were washed first with 500 µL with low detergent buffer (50 mM HEPES pH 7.6, 140 mM NaCl, 0.1% IGEPAL-CA630 (v/v)), followed by three washes, each of 500 µL of wash buffer (50 mM HEPES pH 7.6, 140 mM NaCl). Finally, beads were eluted with mixing in 75 µL of denaturing elution buffer (60 mM Tris pH 6.8, 25% glycerol, 4% SDS(v/v)). Samples from inputs, flowthrough samples, and elution samples were set aside for immunoblotting analysis. Eluates were then snap frozen in liquid nitrogen and stored at -80°C until processed for quantitative mass spectrometry.

### Immuno-blot analysis of immunoprecipitates

Set aside portions of samples from inputs, flowthroughs and elutions from anti-HA or anti-FLAG immunoprecipitation were collected and mixed with 4X sample buffer to a final concentration of 31 mM Tris-HCl pH 6.8, 1% SDS, 12.5% glycerol and 0.005% bromophenol blue before being subjected to gel electrophoresis. Samples were run on 4-20% Criterion TGX Stain free Protein gels (Biorad cat. # 5678095) before transfer to 0.45 µ PVDF membrane (Thermo Fisher Scientific cat. # 88518) via wet transfer. Membranes were blocked with Intercept TBST blocking buffer (Licor cat. # 927-60001) followed by overnight incubation in 5% Bovine serum albumin reconstituted in Tris buffered saline with 0.01% Tween-20 (TBST) with the indicated primary antibodies: anti-FLAG-M2, mouse monoclonal (Sigma, Cat #: F1804, 1:2000), anti-HA c6E2 mouse monoclonal (Cell Signaling Technology, Cat #: 2367, 1:1000); anti-FBXO7 cD8L4E rabbit monoclonal (Cell Signaling Technology, Cat #: 57981, 1:2000); anti-PSMF1 rabbit polyclonal (Proteintech, cat. # F1804; 1:2000); anti-PSMB5 rabbit polyclonal (Bethyl cat Cat #: A303-847A, 1:2000); and anti-CUL1 mouse monoclonal c2H4C9 (ThermoFisher cat # 32-2400. Following washes in TBST, antibodies were detected with 800CW or 680RD conjugated anti-mouse or anti-rabbit antibodies (Licor cat. # 926-32211 and 926-68070) imaged using an ChemiDoc MP Imaging System (Bio-Rad Laboratories cat #. 12003154). When necessary, blots were stripped (Sigma ReBlot Plus Strong antibody stripping solution) prior to re-probing.

### Quantitative mass spectrometry of immunoprecipitates

Elution samples from Immunoprecipitations were reduced with 5 mM TCEP for 15 minutes, alkylated with 10 mM iodoacetamide for 20 minutes in the dark, and quenching the reaction with 15 mM DTT. Following reduction and alkylation, peptide digestion was performed on beads following buffer exchange using SP3 method^76^ adapted for the Kingfisher Flex. Each sample was incubated with 28 µL of a 1:1 mixture carboxylate modified, magnetic SpeedBeads of 1µm average and 0.7-1.10 µm average diameter (Cytiva cat #:65152105050250 and cat #:45152105050250) as a 5% slurry. After incubating these samples for 5 minutes, samples were mixed with an equal volume of pure ethanol, then washed three times, each with 200 µL of 80% ethanol before digestion overnight at 37°C with agitation on a microplate mixer with sequencing grade modified trypsin (Thermo Scientific, cat#: 90059) and Lys-c (Fujifilm Wako cat#: NC9242798). Following digestion, peptide-containing buffer was transferred to a fresh plate for labeling with TMTpro 18 reagents (Thermo), removing the remaining SpeedBeads and incubated for 90 minutes hour at 25°C in a solution containing 30% (v/v) acetonitrile. Following labeling each sample was quenched with 0.5% hydroxylamine (w/v), pooled in equal ratios, diluted to a final concentration of 5% acetonitrile and acetified with formic acid. The pooled sample was desalted using a C18 StageTip^77^, dried in a vacuum centrifuge and resuspending in 5% formic acid, 5% acetonitrile for analysis by liquid chromatography and tandem mass spectrometry (LC-MS). Mass spectrometric data were collected on an Orbitrap Ascend MultiOmics mass spectrometer coupled to a Vanquish Neo UHPLC. Approximately 1µg of peptide was separated at a flow rate of 300 nL/min on a 100 µm capillary column that was packed with 35 cm of Accucore 150 resin (2.6 μm, 150Å; ThermoFisher Scientific). The gradient was 7-25% Buffer B (95% acetonitrile, 0.125% formic acid) which was mixed into buffer A (5% acetonitrile, 0.125% formic acid) over a 135 min gradient.

The scan sequence began with an MS1 spectrum (Orbitrap analysis, resolution 60,000, 350-1350 Th, automatic gain control (AGC) target is set to 100%, maximum injection time set to 50ms, RF lens was set to 30%). The hrMS2 stage consisted of fragmentation by higher energy collisional dissociation (HCD, normalized collision energy 36%) and analysis using the Orbitrap (AGC 200%, maximum injection time 120 ms, isolation window 0.7 Th, resolution 45,000 with TurboTMT activated). Data were acquired using the FAIMSpro interface the dispersion voltage (DV) set to 5,000V, the compensation voltages (CVs) were set at - 30V, -50V, and -70V. The sample was analyzed a second time using a different set of CV (-40V, -60V, and -80V). The number of dependent scans was set to 25 scans per MS1 per CV. Dynamic exclusion list (duration=120s) was shared among CVs.

### Identification and alignment of FBXO7 and PI31 orthologs

To analyze sequence conservation across PI31 and FBXO7 orthologs, the AlphaFold Clusters^78^ containing human PI31 (uniprot ID: Q92530) or human FBXO7 (uniprot ID: Q9Y3I1) were aligned with Clustal Omega (https://www.ebi.ac.uk/jdispatcher/msa/clustalo) and visualized with Consurf (https://consurf.tau.ac.il/) and Protein Domain Designer (https://domaindesigner.farnunglab.com/).

To compare conservation within the FP and PRR domains of FBXO7 and PI31, Clustal Omega alignments of PI31 or FBXO7 sequences were constructed from a diverse set of species with well-annotated proteomes with unambiguous orthologs for FBXO7, PI31, and SKP1, identified with inParanoidDB (https://inparanoidb.sbc.su.se/): *Homo sapiens, Bos taurus, Mus musculus, Gallus gallus, Callorhinchus milii, Xenopus laevis, Danio rerio, Chlorella variabilis, Apostichopus japonicus, Nemastostella vectensis, Daphnia pulex*, and *Oryza sativa*). Alignments of these sequences were mapped onto the FP domains of FBXO7 and PI31 using ChimeraX version 1.10 (ref^79^).

### Cloning, protein expression and purification for biochemical and structural studies

All cDNAs were obtained from an in-house human cDNA library (Max Planck Institute of Biochemistry). Baculovirus transfer vectors for expression of open gate 20S core particles were assembled as previously described^66^. In short, the expression cassettes coding for α3 and α7 in pACEBac1-PSMA1/PSMA2/PSMA3/PSMA4 (Addgene plasmid #217893)^66^ were replaced with expression cassettes lacking residues 2-9 of α3 and residues 2-12 of α7, to obtain the vector pACEBac1-PSMA1/PSMA2/NΔ12-PSMA3/NΔ9-PSMA4 (Addgene plasmid #252020). This vector as donor and pACEBac1-PSMA5/PSMA6/PSMA7 (Addgene plasmid #217894)^66^ as acceptor was used to assemble the vector pACEBac1-PSMA1/PSMA2/NΔ12-PSMA3/NΔ9-PSMA4/PSMA5/PSMA6/PSMA7 (Addgene plasmid #252021) via the MultiBac multiplication module^80^. Expression of open gate 20S core particles was archived by co-infection of insect cells with viruses generated with this vector and the previously published vectors pACEBac1-POMP/PAC1/PAC2/PAC3/PAC4 (Addgene plasmid #214137) and pACEBac1-PSMB1/PSMB2/PSMB3/PSMB4-TEV-2xSTII/PSMB5/PSMB6/PSMB7 (Addgene plasmid #214140)^66^.

Baculovirus transfer vectors for expression of twin-strep-tagged or GST-tagged FBXO7–SKP1complexes, and His8-tagged-PA28αβ (PSME1/PSME2) were generated by insertion of the respective cDNAs into pACEBacDual, which was generated by exchange of the single expression cassette of pACEBac1^80^ with the dual expression cassette of pFBDM^81^. Baculovirus transfer vectors for expression of His8-tagged PI31 were generated by insertion of their respective cDNAs into pACEBac1^80^. For co-expression of FBXO7– SKP1–PI31 complexes PI31 cDNAs were cloned into the donor vectors pIDC^80^, and subsequently fused with the corresponding pACEBacDual-SKP1–FBXO7 acceptor vector. Point mutants were generated by site-directed mutagenesis. Using this approach the following baculovirus transfer vectors were constructed: pACBacDual-PSME2/His8-PP-PSME1 (Addgene plasmid #252037), pACEBacDual-pIDC-SKP1/2xSTII-TEV-FBXO7-169-522/His8-PP-PI31-1-271 (Addgene plasmid #252038), pACEBacDual-pIDC-SKP1/2xSTII-TEV-FBXO7-169-522/His8-PP-PI31-1-151 (Addgene plasmid #252039), pACEBacDual-pIDC-SKP1/2xSTII-TEV-FBXO7-169-398/His8-PP-PI31-1-271 (Addgene plasmid #252040), pACEBacDual-pIDC-SKP1/2xSTII-TEV-FBXO7-169-398/His8-PP-PI31-1-151 (Addgene plasmid #252041), pACEBacDual-SKP1/GST-TEV-FBXO7-169-522 (Addgene plasmid #252042), pACEBacDual-pIDC-SKP1/2xSTII-TEV-FBXO7-169-398/His8-PP-PI31-1-271-L228V (Addgene plasmid #252043), pACEBacDual-pIDC-SKP1/2xSTII-TEV-FBXO7-169-398/His8-PP-PI31-1-271-R242G (Addgene plasmid #252044), pACEBacDual-pIDC-SKP1/2xSTII-TEV-FBXO7-169-398/His8-PP-PI31-1-271-R242C (Addgene plasmid #252045), pACEBacDual-pIDC-SKP1/2xSTII-TEV-FBXO7-169-398/His8-PP-PI31-1-231-R231X (Addgene plasmid #252046).

Sf9 insect cells (Thermo Fisher Scientific) were cultured in serum-free Ex-cell 420 medium (Sigma-Aldrich), and Trichoplusia ni High FiveTM (Thermo Fisher Scientific) were cultured in protein free ESF 921 insect cell culture media (Expression Systems LLC). Bacmid preparations, virus amplification in Sf9, and protein expression in Trichoplusia ni High FiveTM insect cells carried out according to standard protocols^81-83^.

All FBXO7–SKP1 complexes, PI31 constructs, PA28αβ, as well as open gate and wt 20S CPs were first purified by standard affinity purification on either Strep-Tactin (R) Sepharose (IBA), PureCube 100 Ni-INDIGO agarose (Cube Biotech), or Glutathione Sepharose HP (Cytiva), and subsequently by ion exchange chromatography on a 5-ml HiTrap Q HP column (Cytiva). FBXO7–SKP1–PI31complexes were purified by tandem affinity purification on Strep-Tactin (R) Sepharose (IBA), and PureCube 100 Ni-INDIGO agarose (Cube Biotech). As a final step all proteins were purified by size exclusion chromatography on Superdex200 Increase 10/300 columns (Cytiva) or Superose6 Increase 10/300 columns (Cytiva) in buffer A 25 mM HEPES pH 7.5 (KOH), 150 mM NaCl, 0.5 mM TCEP. FBXO7-129-398–SKP1–PI31-1-151 (ref.^19^) and CUL1-NTD^84^ were expressed and purified according to previously described protocols.

### Proteasome activity assays

Proteasome activity assays with recombinant OG-20S CPs and multimeric recombinant FBXO7–SKP1–PI31 complexes were carried out as described previously^56^. Briefly, OG-20S CPs (15 nM) were pre-incubated with the indicated complexes at 37.5-fold molar excess relative to each class of catalytic active sites of the twofold symmetric 20S CP, at 37°C for 15 min and subsequently added to reaction mixtures containing fluorogenic peptides to assay the three distinct catalytic activities of the 20S CP in buffer containing 40 mM HEPES (pH 7.5), 40 mM NaCl, 5 mM MgCl2, and 1 mM DTT. Activity was monitored by cleavage of the following model substrates: β5 chymotrypsin-like activity: Suc-LLVY-AMC (Merck, 539141) at 50 µM, β2 trypsin-like activity: Z-VVR-AMC (Enzo Life Sciences, BML-P199) at 100 µM, and β1 caspase-like activity: Z-LLE-AMC (Merck, 539142) at 50 µM. Fluorescence resulting from AMC release was monitored at Ex/Em = 360/460 nm nm over a time course of 15 min in a multititer plate reader (BMG Labtech, CLARIOstar Plus), and activity was defined as the rate of AMC release. All assays were performed in triplicate and analyzed by one-way ANOVA using GraphPad Prism 9.

Proteasome activity assays with recombinant human OG-20S CPs or native human 20S CP purified from erythrocytes (Enzo Life Sciences, BML-PW8720) and peptides derived from FBXO7 or PI31 were carried out as described previously^51^. Briefly, proteasomes (5 nM) were pre-incubated for 20 min on ice with the indicated peptides at 1 µM or at the indicated concentrations in buffer containing 50 mM Tris-HCl (pH 7.5), 1 mM EDTA, 5 mM MgCl2, 10% glycerol, and 0.02% SDS. Site-specific fluorogenic peptides were used to assay the three distinct proteasome active sites. β5 chymotrypsin-like activity was monitored using Suc-LLVY-AMC (Bachem I-1395) at 100 μM for 45 min at 30°C. β2 trypsin-like activity was monitored using Boc-LRR-AMC (Bachem I-1585) at 300 μM for 90 min at 30°C. β1 caspase-like activity was monitored using Z-LLE-AMC (Bachem I-1945) at 300 μM for 90 min at 30°C. Reactions were quenched by addition of 1% SDS, and fluorescence resulting from AMC release was measured at Ex/Em = 360/460 nm using a VersaFluor fluorometer (Bio-Rad). Fluorescence values were background-subtracted using control reactions lacking 20S CP, and activity was expressed relative to control reactions lacking inhibitors.

FBXO7 peptides were synthesized by WSHT Bio (Shanghai, China) with >95% purity at a 5 mg synthesis scale. All peptides were validated by high-performance liquid chromatography (HPLC) and mass spectrometry. Lyophilized peptides were reconstituted in water and stored at −20°C.

In all assays, the fluorescent substrate was used at a concentration substantially exceeding that of the tested peptides, thereby excluding substrate competition as the primary cause of the observed inhibition.

### Structure determination by transmission electron cryo-microscopy

The sample for the CUL1–FBXO7–SKP1–PI31 structure was prepared by mixing of 20 µL CUL1-NTD at 35 µM and 20 µL FBXO7-129-398–SKP1–PI31-1-151 at 25 µM in buffer A and incubation at 37°C for 15 min. For plunging samples were diluted to a final concentration of 0.3 mg ml^-1^. The sample for the open gate 20S CP–PI31 structure was obtained by co-expression in insect cells and subsequent affinity purification via the twin-strep-tag (20S CP) and His8-tag (PI31). Final buffer exchange to buffer A was carried out with gravity flow PD-10 columns (Cytiva). The sample for the open gate 20S CP–FBXO7–SKP1 structure was obtained by size exclusion chromatography on a Superose6 Increase 10/300 columns (Cytiva) in buffer A. To this end 90 µL open gate 20S CP at 9.5 µM and 210 µL GST-FBXO7-169-522–SKP1 at 185 µM were incubated 30°C for 15 min and gel filtrated. For plunging of OG-20S CP-PI31 and OG-20S CP-FBXO7-SKP1 complexes were concentrated to 0.45 and 0.66 ml^-1^, respectively. For cryo-EM grid preparation Quantifoil R1.2/1.3 Cu-200 grids (Quantifoil Micro Tools GmbH) were glow discharged for 45 seconds in a PDC-32G-2 (Harrick Plasma Inc.), 3.5 µL samples were applied, and plunge frozen in liquid ethane/propane mix with a Vitribot Mark IV (Thermo Fisher Scientific) at 4°C operated at 100% humidity.

All grids were pre-screened for optimal ice thickness and particle distribution on an Glacios or Arctica cryo-TEM (Thermo Fisher Scientific) operated at 200 kV equipped with a K2 Summit (Glacios) or K3 (Arctica) direct electron detector (DED) camera (Gatan). Data collections were set up with SerialEM version 4.1 (ref.^85^) utilizing coma-corrected beam-image shift. Data collection of the CUL1–FBXO7–SKP1–PI31 dataset was carried out on a Glacios cryo-TEM operated at 200 kV equipped with a K2 Summit direct electron detector (DED) camera (Gatan). With a 5x5 multi hole record acquisition scheme one movie per hole was recorded in counting mode at a pixel size of 1.181 Å/pixel with a nominal magnification of 36000x. A total dose of 63.55 e-/Å^2^ was fractionated over 40 frames, with a target defocus range set to -1.0 µm to -2.6 µm. Data collection of the OG-20S CP dataset was carried out on a Arctica cryo-TEM operated at 200 kV equipped with a K3 direct electron detector (DED) camera (Gatan). With a 5x5 multi hole record acquisition scheme one movie per hole was recorded in counting mode at a pixel size of 0.919 Å/pixel with a nominal magnification of 36000x. A total dose of 63.0 e-/Å^2^ was fractionated over 40 frames, with a target defocus range set to -0.8 µm to -2.1 µm. Data collection of the PI31-bound and FBXO7– SKP1-bound open gate 20S CP datasets was carried out on a Titan Krios G2 cryo-TEM (Thermo Fisher Scientific) operated at 300 kV equipped with a Bio Quantum post-column energy filter (Gatan, 10eV) and K3 direct electron detector (DED) camera (Gatan). With a 5x5 multi hole record acquisition scheme three movies per hole were recorded at a pixel size of 0.8512 Å/pixel with a nominal magnification of 105000x in counting mode (no CDS). A total dose of 63.0 e-/Å^2^ (PI31-bound OG-20S CP) or 45.0 e-/Å^2^ (FBXO7-bound OG-20S CP) was fractionated over 30 frames, with a target defocus range of -0.6 µm to -2.2 µm (for details see **Extended Data Table 1**).

Data processing was performed as follows. For the open gate 20S CP-PI31 dataset movies were on the fly motion corrected with FOCUS^86^, and motion corrected micrographs were imported into cryoSPARC version 4.7.1 (ref.^87^), all subsequent processing steps were carried out with cryoSPARC, for details see **Extended Data Fig. 7e-h**. For the open gate 20S CP-FBXO7-SKP1 and the CUL1-NTD-FBXO7–SKP1–PI31 data sets raw movies were imported into cryoSPARC and all processing steps were carried out with cryoSPARC, for details, see **Extended Data Fig. 7a-f**.

Model fitting was performed as follows. The AlphaFold3 (ref.^61,62^) model of CUL1-1-212–FBXO7-169-398–SKP1–PI31-1-151 were generated with the AlphaFold3 web browser. Regions of Cul1 modelled with low confidence (residues 1-14 and 58-79) were excluded from the model and chains were individually fitted into the cryo-EM map utilizing UCSF ChimeraX version 1.10 (**Extended Data Fig. 4a**).

Model building and refinement was performed as follows. For the open-gate 20S CP-PI31 structure the chains from the published models of the human 20S CP (PDB 8QYO) along with the chains of PI31 (PDB 8FZ6) were individually fitted into the cryo-EM map with ChimeraX. The atomic model for the FBXO7 chains in the OG-20S CP–FBXO7–SKP1 structure as well as unmodeled regions of the PI31 chains in the OG-20S CP–PI31 structure were built *de novo* in Coot version 0.9.8.8 (ref.^88^). Subsequently all models were iteratively refined with Coot and Phenix version 1.21.1 (ref.^89^). For visualization figures were generated with UCSF ChimeraX, and Adobe Illustrator 2026.

## Data availability

Cryo-EM maps and corresponding atomic coordinates have been deposited in the Electron Microscopy Data Bank (EMDB) and the Protein Data Bank (PDB), respectively. Accession codes: EMD-57672 (human SCF complex containing SKP1, CUL1 N-terminal domain, FBXO7 residues 129-398, and PI31 residues 1-151); EMD-57772; PDB: 30HM (human 20S proteasome core particle with α3 and α7 N-terminal deletion); EMD-57773, PDB: 30HN (human 20S proteasome core particle with α3 and α7 N-terminal deletion in complex with PI31); EMD-57774; PDB: 30HO (human 20S proteasome core particle with α3 and α7 N-terminal deletion in complex with FBXO7). Raw cryo-EM movie frames of all datasets have been deposited in the Electron Microscopy Public Image Archive (EMPIAR) under the following accession codes: EMPIAR-XXXXXX, EMPIAR-XXXXXX, EMPIAR-XXXXXX, and EMPIAR-XXXXXX.

Proteomic data associated (.RAW files) are available via ProteomeXchange for immunoprecipitations of FBXO7 with PI31 truncations (PXD077028), immunoprecipitations of FBXO7 truncations (PXD077448) and immunoprecipitations of PI31 variants (PXD077016).

Vectors for protein expression were submitted to Addgene and a comprehensive list of these constructs, datasets, and other resources associated are provided in the associated key resource table (https://dx.doi.org/10.5281/zenodo.20276561).

Detailed protocols associated with this work are available on protocols.io (https://dx.doi.org/10.17504/protocols.io.261geymowv47/v1;

https://dx.doi.org/10.17504/protocols.io.3byl4p4m8lo5/v1;

https://dx.doi.org/10.17504/protocols.io.81wgbj3rovpk/v1;

https://dx.doi.org/10.17504/protocols.io.6qpvrb562lmk/v1;

https://dx.doi.org/10.17504/protocols.io.q26g7o2m9vwz/v1;

https://dx.doi.org/10.17504/protocols.io.eq2lyok6wgx9/v1).

### Acknowledgments

We thank D. Bollschweiler and T. Schäfer for access to the MPIB cryo-EM facility and their assistance with data collection. We also acknowledge J. Rajan Prabu for maintaining processing infrastructure. This work was supported by Aligning Science Across Parkinson’s (ASAP, grants 000282, 024268, and 025160), the Max Planck Society, and the Leibniz Prize of the Deutsche Forschungsgemeinschaft (DFG, German Research

Foundation: SCHU 3196/1-1 to B.A.S.), as well as by National Institutes of Health (NIH) grants R01-GM144367 (to J.H.), R01-AG011085, and R01-NS083524 (to J.W.H.), and R35-GM156406 (to J.A.P.).E.A.G. was supported by a postdoctoral fellowship from the Edward R. and Anne G. Lefler Center for the Study of Neurodegenerative Disorders. The funders had no role in study design, data collection and analysis, decision to publish or preparation of the manuscript. For the purpose of open access, the authors have applied a CC-BY public copyright license to the author-accepted manuscript version arising from this submission.

## Author information

F.A., E.A.G., J.W.H. and B.A.S. designed the research;F.A., E.A.G., E.S. and B.S.M. contributed new reagents;F.A., E.A.G., E.S, J.A.P., E.F.P, D.F., and J.H. performed the research; F.A., E.A.G. E.S. analyzed the data; J.W.H., J.H. and B.A.S. acquired funding; F.A., E.A.G., J.H., J.W.H. and B.A.S. wrote the manuscript with input from all authors.

---

**Extended Data Table 1:**
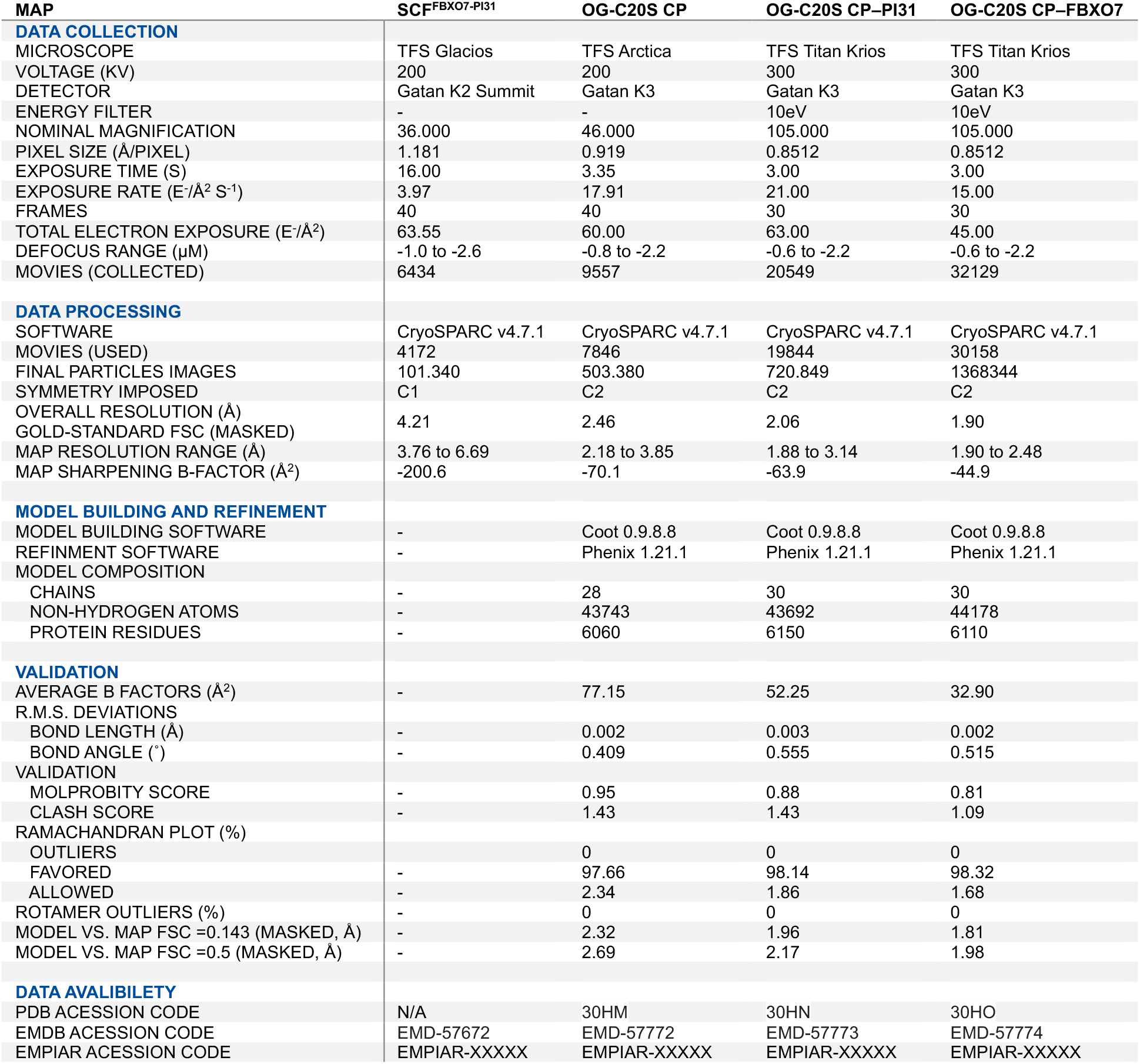
Cryo-EM data collection, refinement and validation statistics.

**Extended Data Fig. 1.**
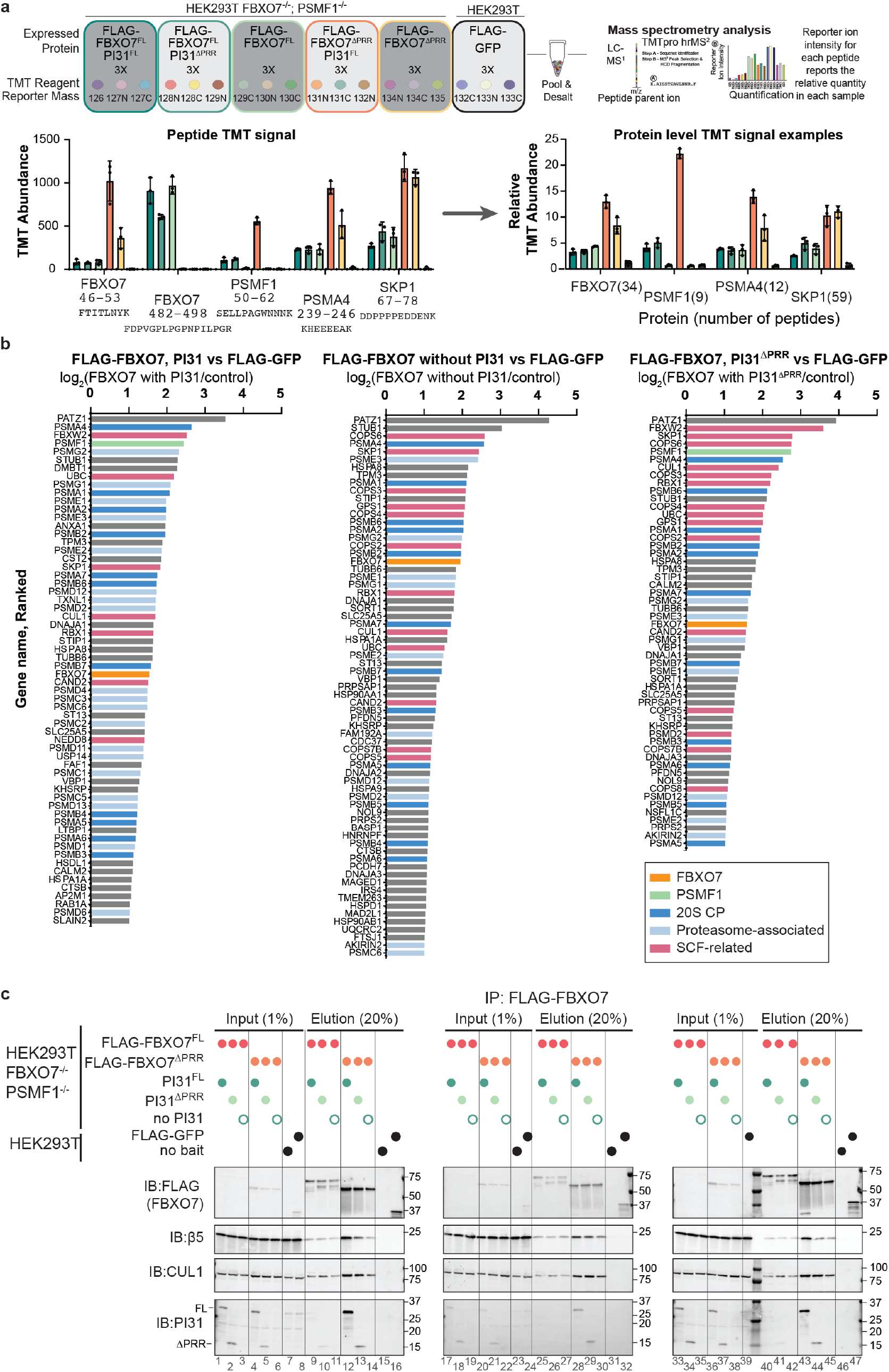
Proteomic analysis of FBXO7 interacting proteins dependence on PI31 and PI31 PRR. (**a**) Experimental design for analysis of FBXO7 interacting proteins. The indicated FBXO7 or PI31 proteins were stably expressed in HEK293T cells lacking both FBXO7 and PI31 and triplicate cultures used for α-FLAG immunoprecipitation and TMT-proteomics using the indicated TMT reporter ion reagents. Individual peptides with corresponding TMT-reporter intensities are mapped to each protein, reporting abundance. Parental HEK293T cell lysates containing added FLAG-GFP were used as a negative control. (**b**) Proteins enriched in FLAG-FBXO7 immunoprecipitates compared to control plotted by ranked log2FC. Color code: pink: SCF components, COP9 signalosome, and CAND exchange factor; dark blue: 20S CP subunits; light blue: 19S RP subunits and binding proteins, alternative caps and proteasome assembly chaperones; gray, non-proteasome or SCF-related proteins. (**c**) Immunoblots of triplicate samples from the experiment outlined in panel **a** and in **Fig. 1 b–d**. The indicated amounts of input and eluate from α-FLAG immunoprecipitates are shown. Immunoblots were probed with the indicated antibodies.

**Extended Data Fig. 2.**
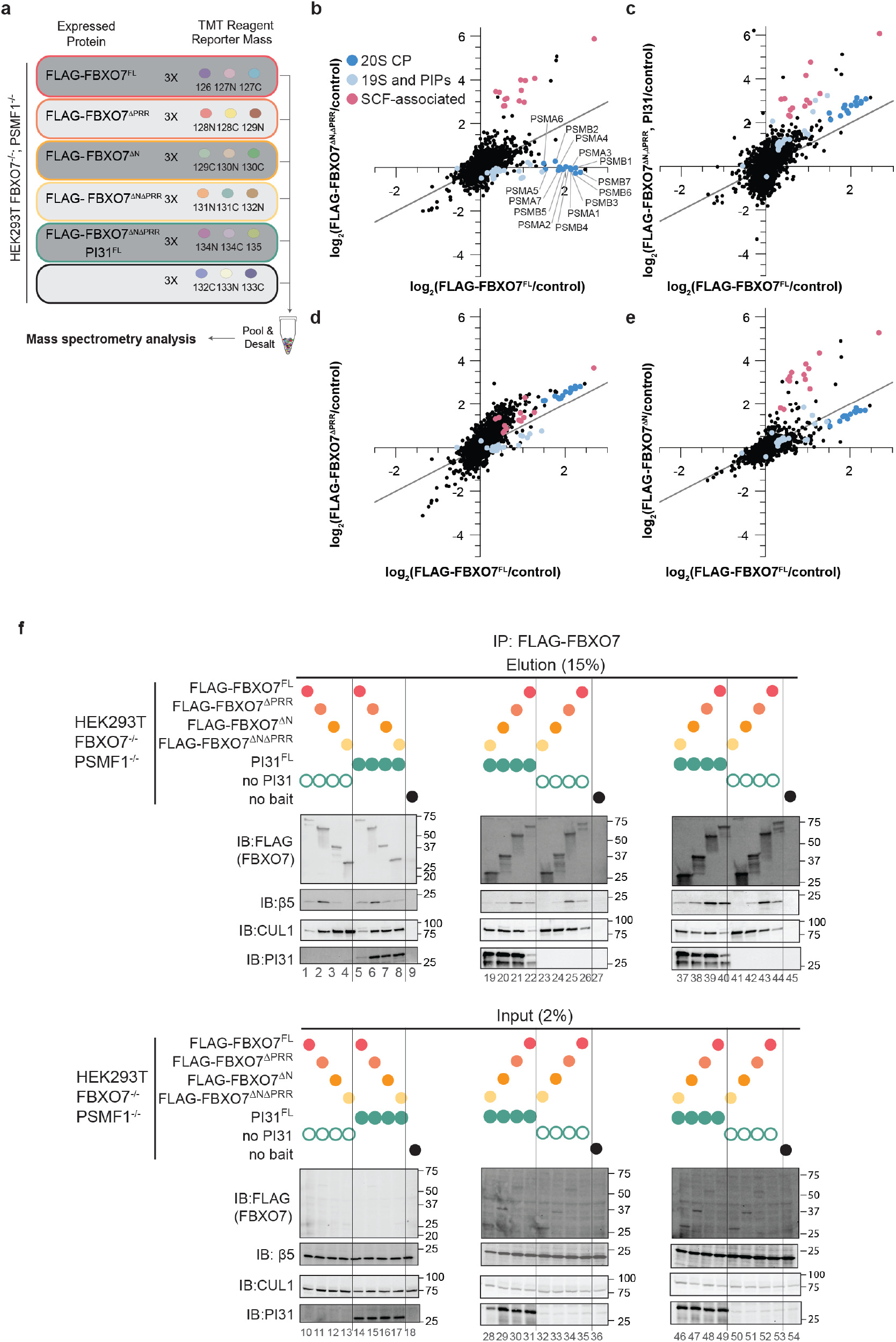
Analysis of FBXO7 domains impact on interacting proteins. (**a**) Experimental design for analysis of FBXO7 interacting proteins. The indicated FBXO7 or PI31 proteins were stably expressed in HEK293T cells lacking both FBXO7 and PI31 with triplicate cultures used for α-FLAG immunoprecipitation and TMT-proteomics using the indicated TMT reporter ion reagents. (**b**) Correlation plots of log2FC (versus control) for enriched proteins from FLAG-FBXO7^FL^ (x-axis) and FLAG-FBXO7^ΔNΔPRR^ (y-axis) immunoprecipitates. All 20S CP subunits are labeled by name. 20S CP components (dark blue) SCF components, COP9 signalosome, and CAND exchange factors (pink), and proteasome-related components including 19S RP subunits and interacting proteins, alternative proteasome caps, and proteasome assembly factors (light blue) are highlighted by color. (**c**) Correlation plots of log2FC (versus control) for enriched proteins from FLAG-FBXO7^FL^ (x-axis) and FLAG-FBXO7^ΔNΔPRR^ (+PI31) (y-axis) immunoprecipitates. Color coding is as in panel **b**. (**d**) Correlation plots of log2FC (versus control) for enriched proteins from FLAG-FBXO7^FL^ (x-axis) and FLAG-FBXO7^ΔPRR^ (y-axis) immunoprecipitates. Color coding is as in panel **b**. (**e**) Correlation plots of log2FC (versus control) for enriched proteins from FLAG-FBXO7^FL^ (x-axis) and FLAG-FBXO7^ΔN^ (y-axis) immunoprecipitates. Color coding is as in panel **b**. (**f**) Immunoblots of triplicate samples from the experiment outlined in **Extended Data Fig. 1a**. The indicated amounts of input and eluate from α-FLAG immunoprecipitates are shown. Immunoblots were probed with the indicated antibodies.

**Extended Data Fig. 3.**
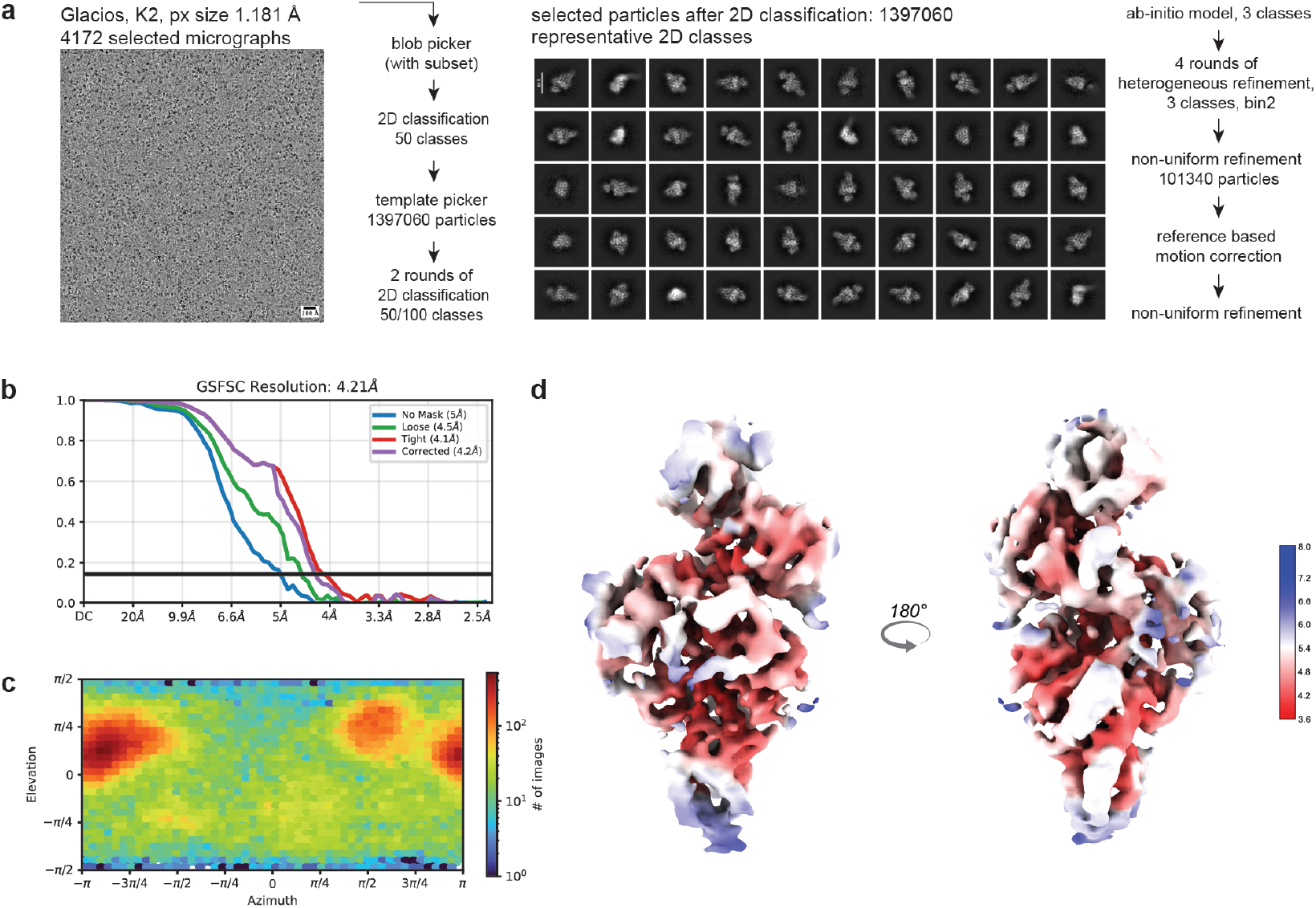
SPA cryo-EM data processing workflow of the FBXO7–SKP1–PI31–Clu1 N-terminal domain complex. (**a**) Data processing scheme with representative micrograph and selected 2D class averages. (**b**) Angular distribution (viewing direction distribution) of particles used for reconstruction, (**c**) Gold-standard Fourier shell correlation (GSFSC) curve after FSC mask auto-tightening, and (**d**) local resolution estimation of the final cryo-EM map reconstruction.

**Extended Data Fig. 4:**
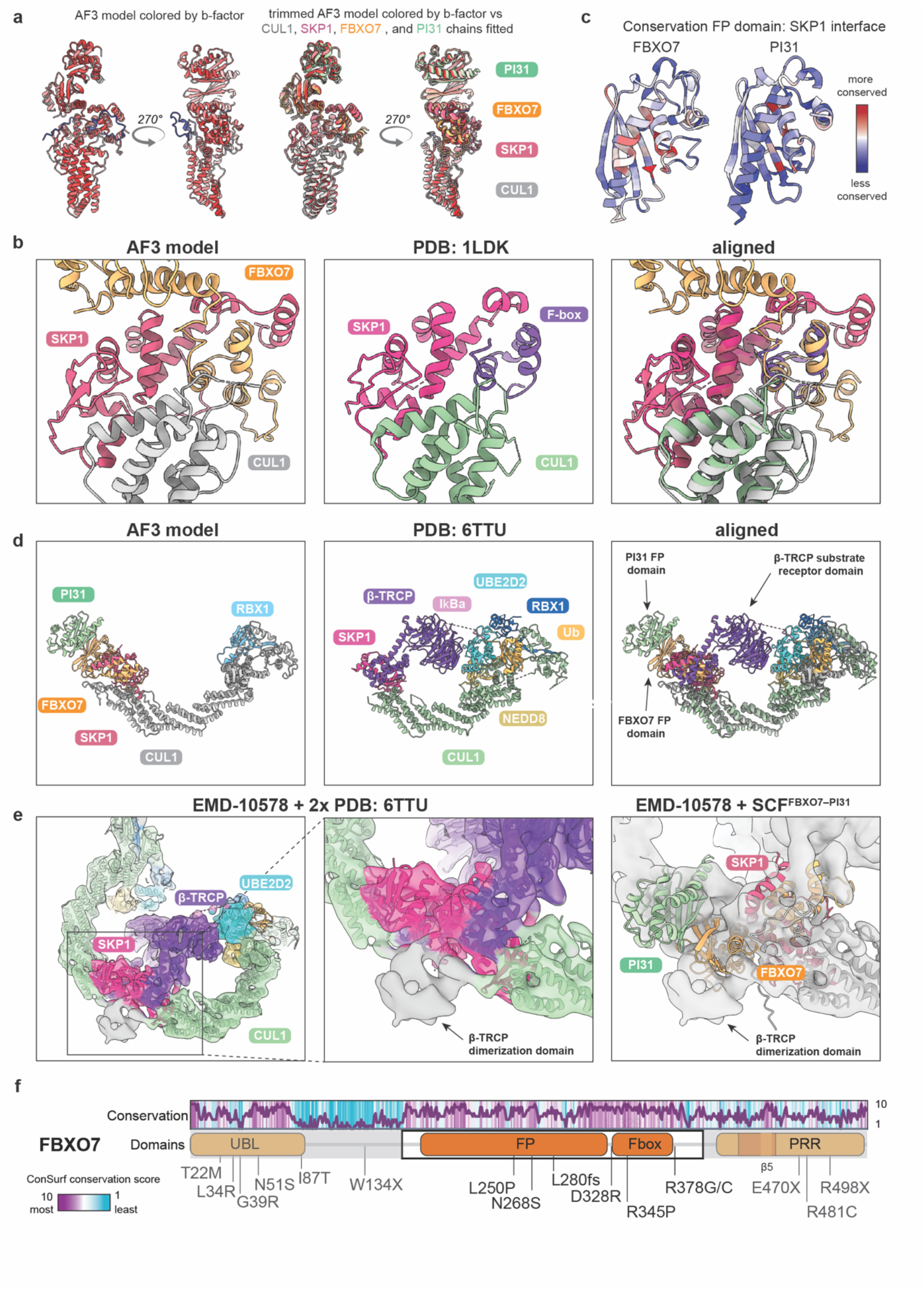
Structural analysis of the SCF^FBXO7–PI31^ complex. **(a)** Ribbon representation of the AlphaFold3 (AF3) model of the SCF^FBXO7–PI31^ complex colored by predicted confidence (B-factor) shown from two orientations (left panel). Trimmed AF3 model after removal of low-confidence regions overlayed with individually fitted chains into the cryo-EM density, colored as in Fig. 2, with PI31 in seafoam green, FBXO7 in orange, SKP1 in pink, and CUL1 in gray shown from two orientations (right panel). **(b)** Comparison of the SCF^FBXO7–PI31^ complex with the published canonical SCF complex structure (PDB: 1LDK). AF3 model of the SCF^FBXO7–PI31^ complex (left panel), canonical SCF complex from PDB 1LDK (middle panel), and aligned structures (right panel), revealing a conserved overall organization of the CUL1–SKP1–F-box architecture are shown. **(c)** Conservation analysis of the FP domains of FBXO7 and PI31 across species. Primary sequence conservation is mapped onto the structures and colored from red (most conserved) to blue (least conserved) highlighting strong conservation within FBXO7 helix 4 (H4). (**d**) Comparison of AF3 model of the SCF^FBXO7–PI31^ complex containing full-length CUL1 (gray) and RBX1 (light blue) (left panel) with a trapped ubiquitin-transfer intermediate structure (PDB: 6TTU) (middle panel). In the published structure, ubiquitin (Ub; yellow) linked to the E2 enzyme UBE2D2 (cyan) is positioned for transfer to the substrate IκBα (light pink) bound by the substrate receptor β-TRCP (purple). The right panel shows aligned structures from the left and middle panels. (**e**) Comparison with the published dimeric cryo-EM structure (EMD-10578) containing CUL1, RBX1, SKP1, β-TRCP including its dimerization domain, UBE2D2, ubiquitin, NEDD8, and IκBα. Overview of the cryo-EM density with two copies of the PDB 6TTU model docked into the map (left panel), close-up views of the β-TRCP dimerization domain with docked PDB 6TTU structures (middle panel), and cryo-EM density with docked model of the SCF^FBXO7–PI31^ complex (right panel) are shown, revealing that the FBXO7 FP and PI31 FP domains partially overlap with the position occupied by the β-TRCP dimerization domain rather than projecting toward the CUL1 C-terminal domain. (**f**) Schematic domain organization of FBXO7. ConSurf conservation scores are mapped onto the primary sequence and colored from most conserved (purple) to least conserved (cyan). Known PPS-associated FBXO7 mutations are indicated.

**Extended Data Fig. 5.**
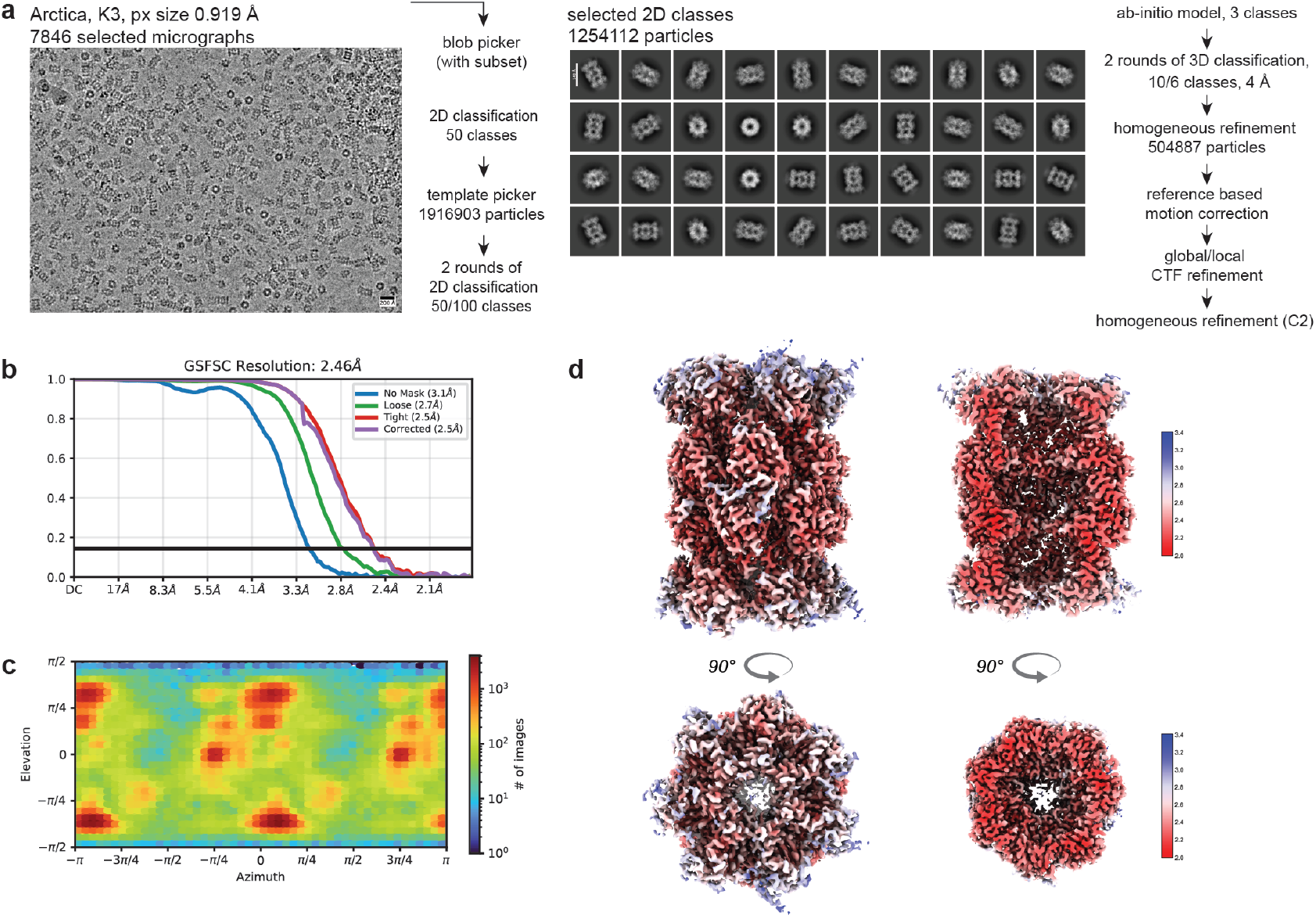
SPA cryo-EM data processing workflow of the open-gate (OG) 20S proteasome core particle. (**a**) Data processing scheme with representative micrograph and selected 2D class averages. (**b**) Angular distribution (viewing direction distribution) of particles used for reconstruction, (**c**) Gold-standard Fourier shell correlation (GSFSC) curve after FSC mask auto-tightening, and (**d**) local resolution estimation of the final cryo-EM reconstruction.

**Extended Data Fig. 6:**
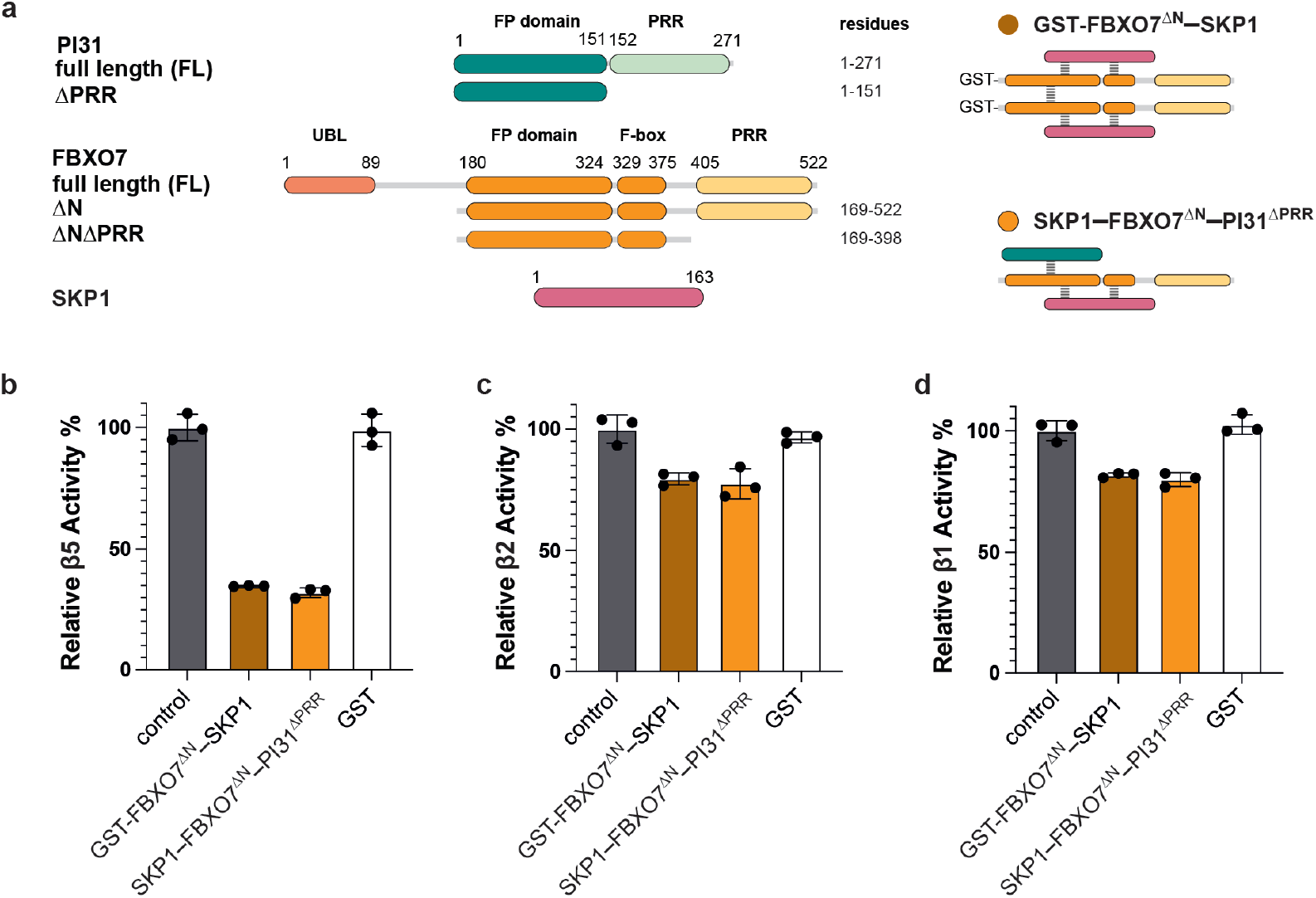
Comparison of proteasome inhibition by complexes containing one or two FBXO7 PRR domains. (**a**) Domain organization of PI31, FBXO7, and SKP1 (left panel), and schematic representation of the composition of homodimeric SKP1– GST–FBXO7ΔN complexes containing two FBXO7 PRRs (“double tail”) and heterotrimeric SKP1–FBXO7ΔN–PI31ΔPRR complexes containing one FBXO7 PRR (“single tail”) (right panel). (**b–d**) Peptidase activity assays with site-specific fluorogenic peptides for (**b**) β5 chymotrypsin-like activity: suc-LLVY-AMC, (**c**) β2 trypsin-like activity: Z-VVR-AMC, and (**d**) β1 caspase-like activity: Z-LLE-AMC. Protease activity of OG-20S CP (15 nM) was assayed in the absence (control) or presence of SKP1–GST-FBXO7^ΔN^ (double tail), SKP1–FBXO7^ΔN^–PI31^ΔPRR^ (single tail), or GST alone (no tail) at 37.5-fold molar excess relative to each class of catalytic active sites. Proteasome activity was normalized to OG-20S CP alone. Mean activity slopes are plotted, with individual replicates shown as dots. Error bars represent standard deviation (n = 3).

**Extended Data Fig. 7.**
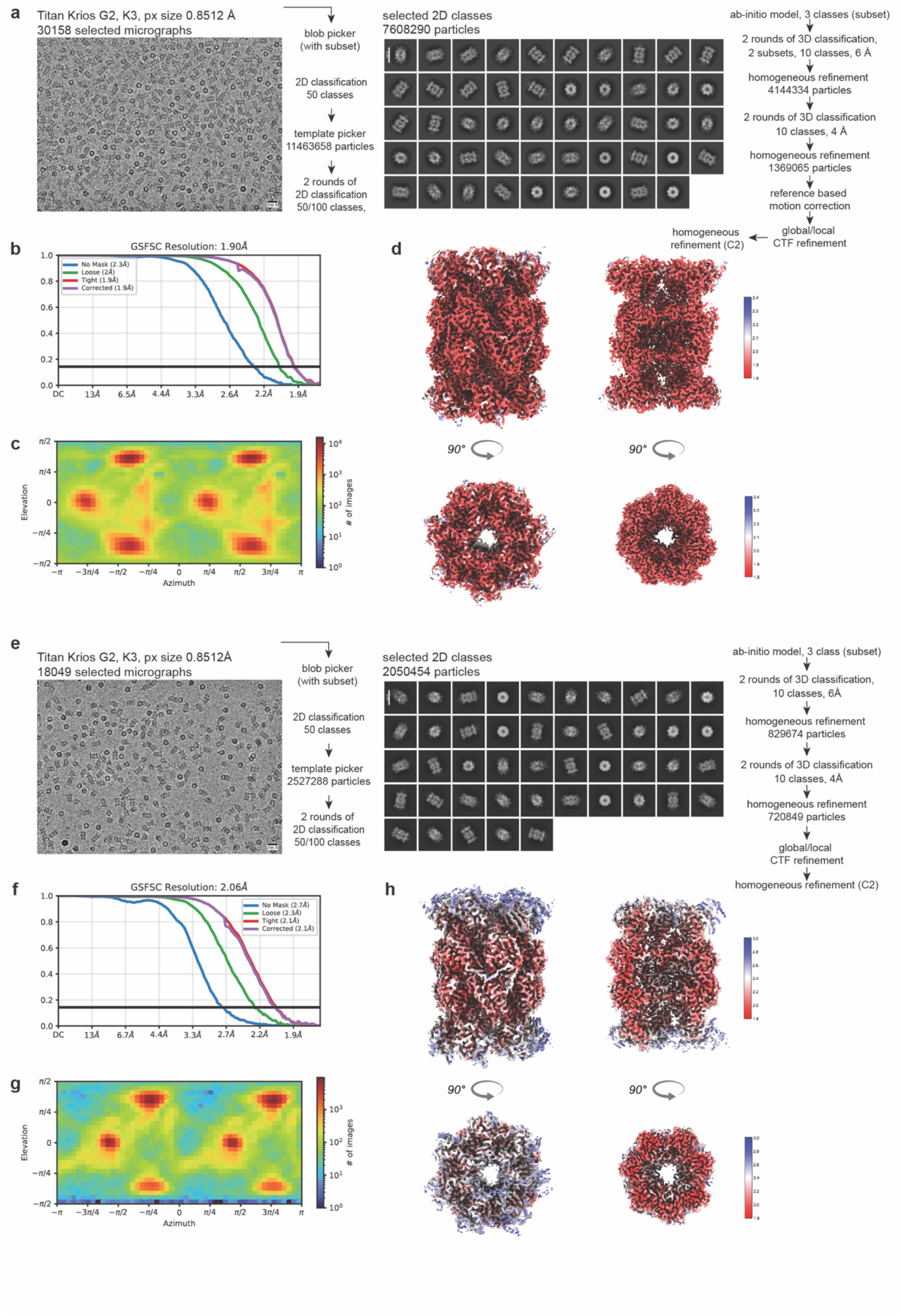
SPA cryo-EM data processing workflow of the FBXO7-OG 20S CP and PI31-OG 20S CP complexes. (**a–d**) FBXO7–20S CP map. (**a**) Data processing scheme with representative micrograph and selected 2D class averages. (**b**) Angular distribution (viewing direction distribution) of particles used for reconstruction, (**c**) Gold-standard Fourier shell correlation (GSFSC) curve after FSC mask auto-tightening, and (**d**) local resolution estimation of the final cryo-EM reconstruction. (**e–h**) PI31–20S CP map. (**e**) Data processing scheme with representative micrograph and selected 2D class averages. (**f**) Angular distribution (viewing direction distribution) of particles used for reconstruction, (**g**) Gold-standard Fourier shell correlation (GSFSC) curve after FSC mask auto-tightening, and (**h**) local resolution estimation of the final cryo-EM reconstruction.

**Extended Data Fig. 8.**
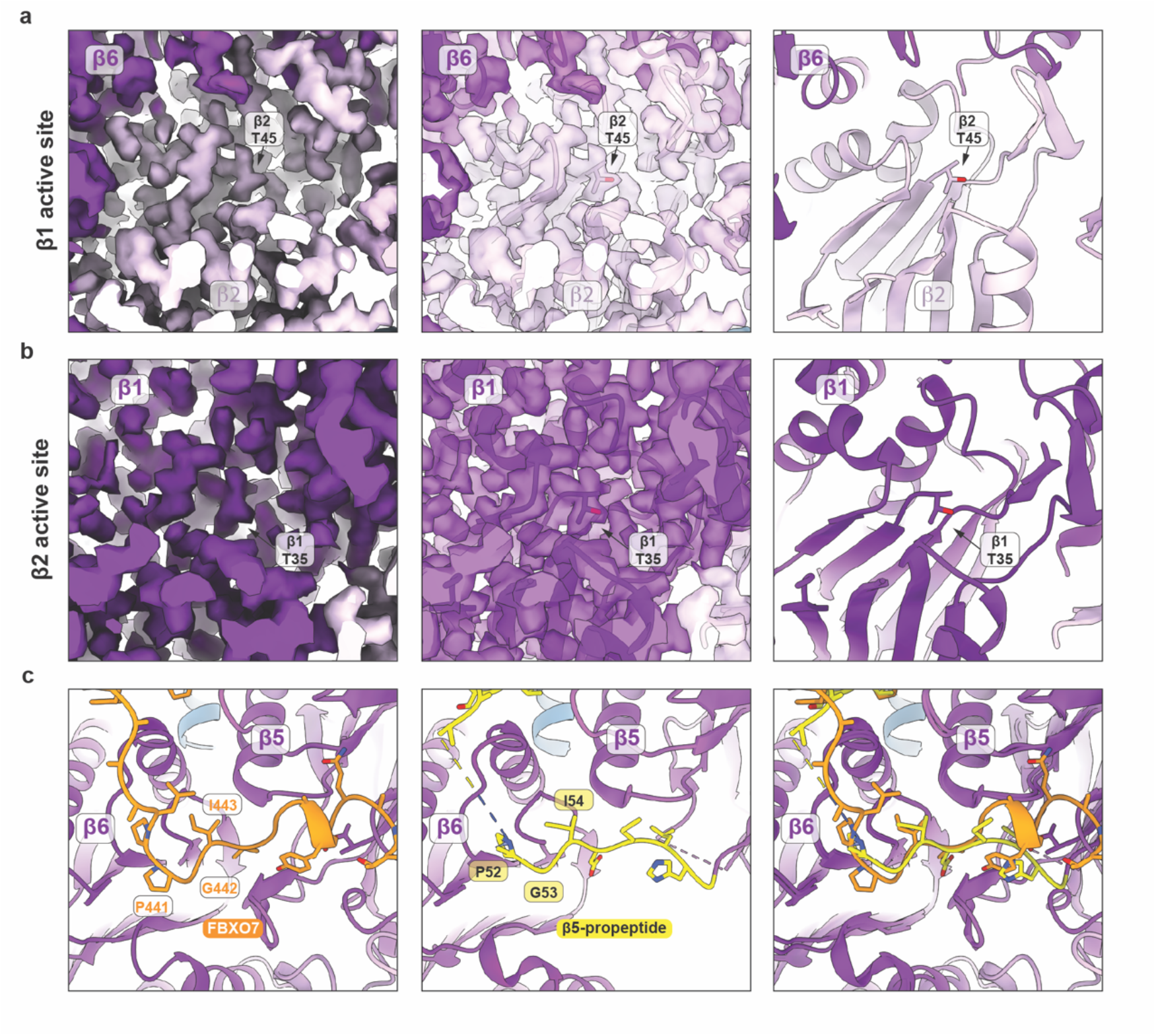
Structural analysis of the FBXO7-bound OG-20S CP complex. **(a–b)** Inset views of the β2 (a) and β1 (b) active sites of the FBXO7-bound OG-20S CP complex. Cryo-EM density (left panels), transparent cryo-EM density with fitted atomic model (middle panels), and atomic model alone (right panels) are shown. Catalytic threonine residues are highlighted. No additional density corresponding to FBXO7 was observed at the β2 or β1 active sites. **(c)** Inset views of atomic models of the β5 active site highlighting a conserved Pro–Gly–Ile motif in FBXO7 (left panel; this study) and the β5 propeptide of prefusion assembly intermediates (middle panel; PDB: 8QYN) occupying similar orientations. The right panel shows an overlay of both structures.

**Extended Data Fig. 9.**
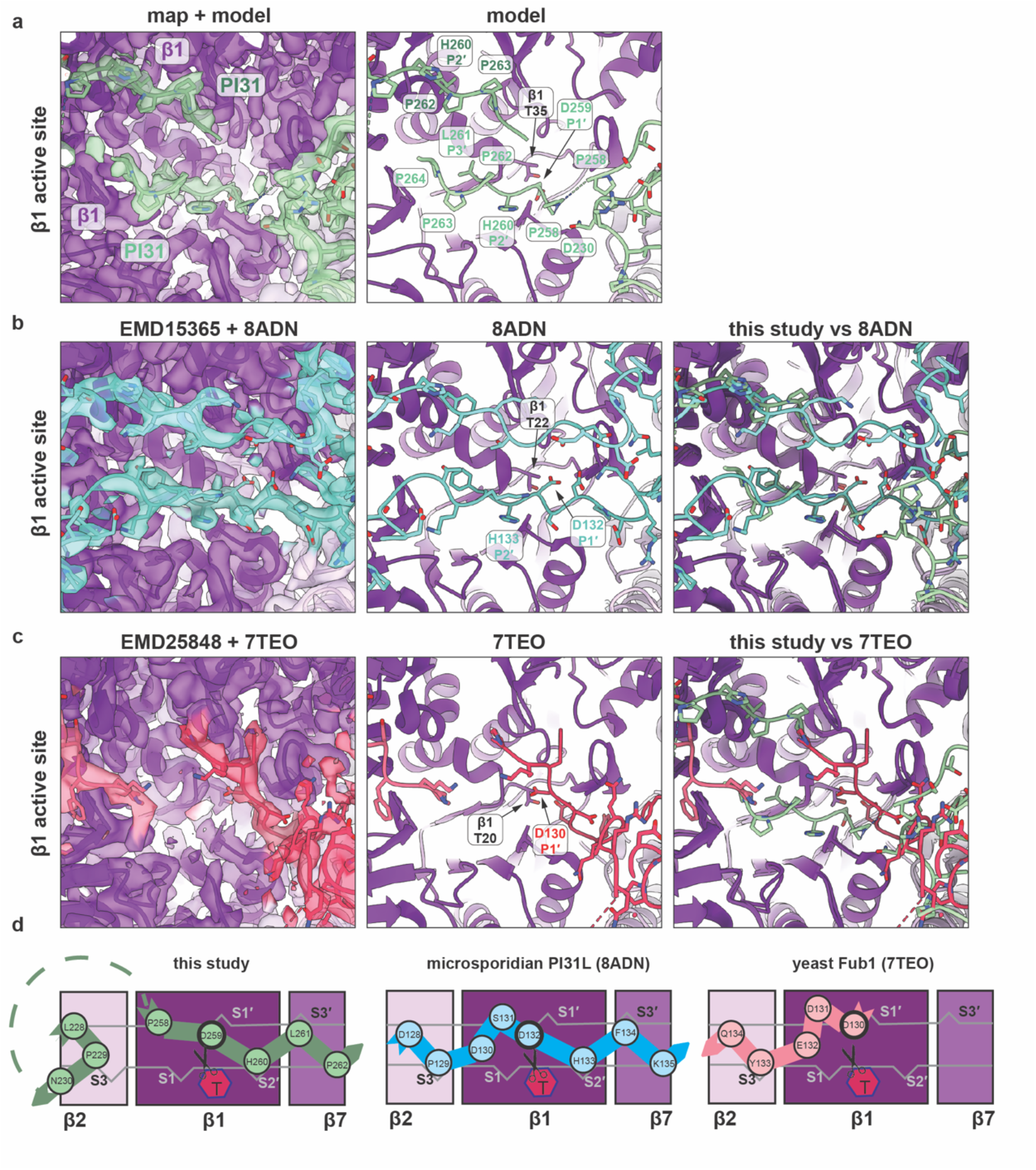
Comparison of the structural mechanism of β1 inhibition by human PI31, microsporidian PI31-like, and their yeast homologue Fub1. (**a**) Human PI31 bound at the β1 site of human OG-20S CP (this study). (**b**) Microsporidian PI31-like bound at the β1 site of microsporidian 20S CP (EMD15365, PDB 8ADN), (**c**) Yeast Fub1 bound at the β1 site of yeast Δα3-20S CP (EMD25848, PDB 7TEO). Transparent cryo-EM densities with fitted atomic model (left panels), atomic models (middle panels) are shown. For comparison overlays of atomic models in (b) and (c) with the atomic model of human PI31 bound at β1 active site (this study) are shown in the right panels. (**d**) Schematic representation of similar but distinct mechanism how human PI31 (left panel), microsporidian PI31-like (middle panel), and yeast Fub1 engage and inhibit the β1 active sites through substrate-mimetic binding.

**Extended Data Fig. 10:**
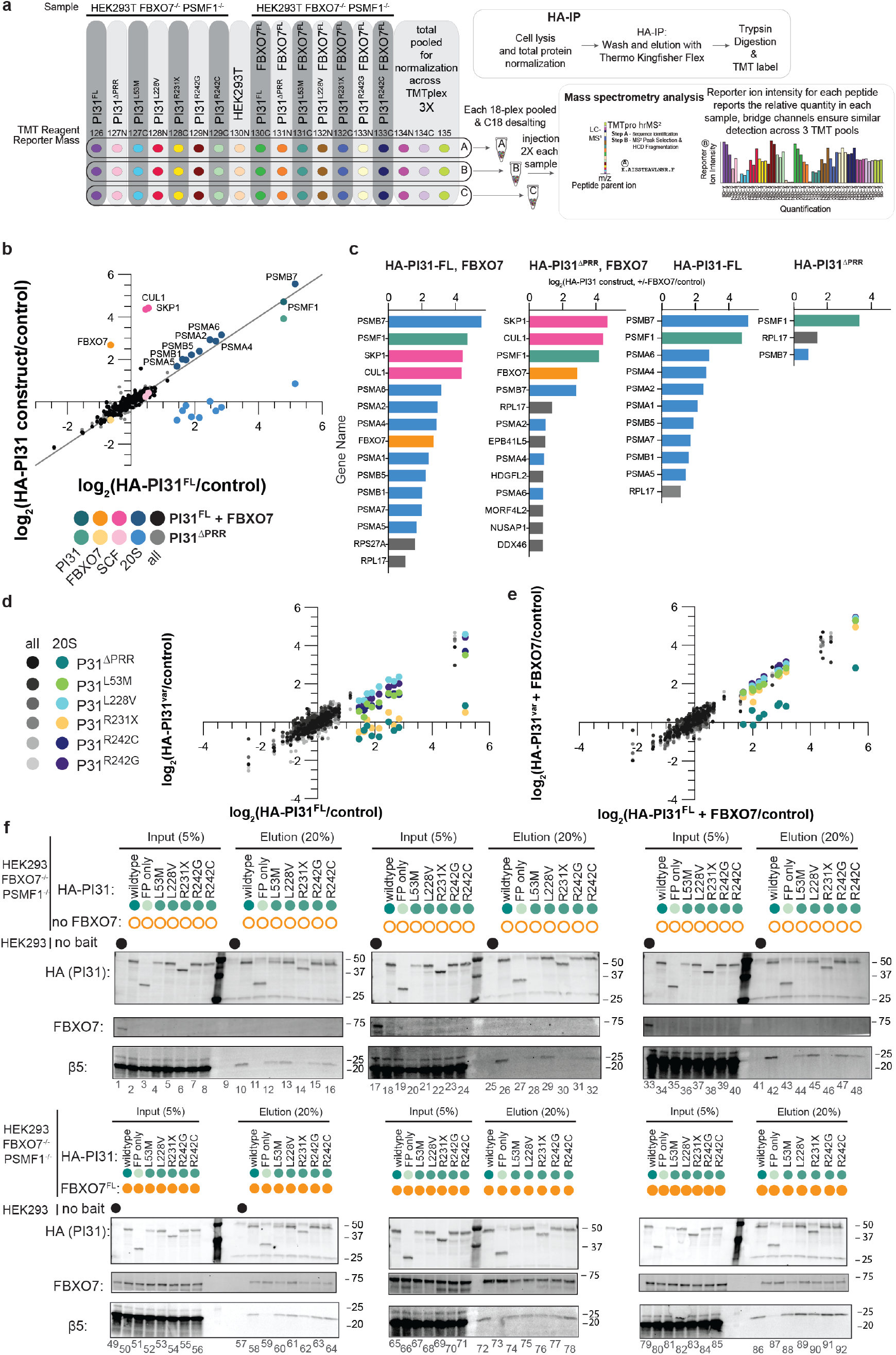
Proteomic analysis of PD-associated PI31 variant interacting proteins and their dependence on FBXO7. (**a)** Experimental design for analysis of PI31 interacting proteins. The indicated FBXO7 or PI31 proteins were stably expressed in HEK293T cells lacking both FBXO7 and PI31 and triplicate cultures used for α-HA immunoprecipitation and TMT-proteomics using the indicated TMT reporter ion reagents. Individual peptides with corresponding TMT-reporter intensities are mapped to each protein, reporting abundance. Parental HEK293T cell lysates were used as a negative control. (**b**) Correlation plots of log2FC (versus control) for enriched proteins from HA-PI31^ΔPRR^ (x-axis) and HA-PI31^FL^ + FBXO7(y-axis). PI31 (seafoam), FBXO7 (orange), SKP1 and CUL1 (pink), and 20S CP subunits (blue) are highlighted and labeled, with all other proteins (grey or black) with at least three peptides in at least two of three replicate samples plotted. (**c**) Proteins with at least 1.75-fold enrichment in HA-PI31 immunoprecipitations of PI31FL with FBXO7, PI31^Δ^PRR with FBXO7 or without FBXO7 listed ranked by enrichment as compared to control. Bars are color coded highlighting PI31 (seafoam), FBXO7 (orange), SKP1 and CUL1 (pink), and 20S proteasome subunits. Correlation plots of log2FC for enriched proteins in HA-PI31 complexes versus control immune complexes from the PI31^ΔPRR^ or the indicated PD-variants of HA-PI31. 20S subunits are colored corresponding to the PI31 variant with all other proteins with at least two peptides detected in at least two samples shown in shades of gray. Immunoprecipitations without FBXO7 expressed shown in (**d**) and the same comparisons for samples with FBXO7 co-expressed in (**e**). (**f**) Immunoblots of triplicate samples from the experiment outlined in panel **a** and in **Fig7. b** and **c**. The indicated amounts of input and eluate from α-FLAG immunoprecipitates are shown. Immunoblots were probed with the indicated antibodies.

